# Lipid mediated ER-stress contributes to the pathogenesis of mitochondrial myopathies

**DOI:** 10.64898/2026.06.10.731336

**Authors:** Inaya Laubach, Guido Primiano, Nneka Southwell, Nicola Rizzardi, Belem Yoval-Sánchez, Christian Bergamini, Serenella Servidei, Alexander Galkin, Giovanni Manfredi, Qiuying Chen, Marilena D’Aurelio

## Abstract

Mitochondrial diseases are a heterogeneous group of genetic disorders caused by impaired oxidative phosphorylation (OxPhos). When skeletal muscle is predominantly affected, they are defined as primary mitochondrial myopathies. Although the genetic causes of mitochondrial myopathies and the resulting bioenergetic impairments are well established, the metabolic drivers behind progressive muscle dysfunction remain poorly defined. This gap in knowledge may contribute to the lack of effective treatments for these disorders. OxPhos defective muscle initiates a metabolic response coordinated by systemic signals which invokes the mobilization of fatty acids from white adipose tissue despite muscle inability to fully oxidize fatty acids due to OxPhos impairment. Here, we show that in human patients with mitochondrial disease and mice with OxPhos defective muscle, increased fatty acid mobilization from white adipose tissue leads to ectopic lipid accumulation and lipotoxicity in skeletal muscle. We find an increase in very long chain ceramides which are mechanistically linked to chronic ER stress, phosphorylation of eIF2α, and activation of ATF4 signaling. We propose that Ph-eIF2α-mediated attenuation of global protein synthesis and ATF4-initiated atrophy pathways contribute to muscle wasting and weakness. Importantly, inhibition of *de novo* synthesis of ceramides with myriocin reduces ER stress and improves muscle proteostasis, body weight, and motor functions in a conditional COX10 KO mouse model of mitochondrial myopathy. Together, these findings highlight altered lipid metabolism as a contributor of mitochondrial myopathy pathogenesis and identify lipid-mediated stress pathways that can be targeted therapeutically.

## INTRODUCTION

Mitochondrial diseases are multi-organ disorders caused by mutations in either mitochondrial (mtDNA) or nuclear (nDNA) genomes, resulting in the impairment of the oxidative phosphorylation (OxPhos) system. Mitochondrial diseases mostly affect tissues with high energetic demands, such as heart, brain, and skeletal muscle [1–3]. When myopathy is the predominant manifestation, they are defined as primary mitochondrial myopathies (PMM). Limited treatments are available for PMM [4], except for nucleosides for specific diseases of mtDNA maintenance [5], and clinical trials lack consensus on metabolic biomarkers or outcome measures [6].

An incomplete understanding of the mechanisms whereby OxPhos defective muscle undergoes metabolic remodeling hinders the development of effective therapies. Clinical symptoms of OxPhos defects include fatigue, muscle weakness, and muscle atrophy. Moreover, PMM patients are often underweight [7] or even cachectic [8], indicating that muscle wasting and decreased adiposity are pathological features of mitochondrial diseases [9]. We and others have identified a coordinated multi-organ metabolic response to OxPhos defective muscle [10, 11]. This metabolic rearrangement initiates in muscle and propagates systemically through myokines, leptin, and glucocorticoid signaling, ultimately resulting in adipose stores depletion, lipid dyshomeostasis, and muscle lipid accumulation.

Adipose tissue plays a major role in the regulation of energy balance and nutritional homeostasis, serving as the main site of energy storage in the form of triacylglycerides (TAG) and as a source of circulating free fatty acids (FA). FA mobilized from white adipose tissue (WAT) are taken up and oxidized by muscle for energy generation. However, when mitochondrial FA β-oxidation (FAO) is impaired, partially oxidized FA derivatives build up in muscle. In accord, elevated levels of FAO intermediates are detected in plasma of Myoclonus Epilepsy and Ragged Red Fibers (MERRF) [11] and Mitochondrial Encephalomyopathy, Lactic Acidosis, and Stroke-like episodes (MELAS) patients, and they are considered validated plasma biomarkers of disease severity [12].

The ectopic storage of lipids in muscle can have toxic effects leading to muscle atrophy, impaired regeneration, sarcopenia [13], and severe myopathy in humans and mice [14, 15]. Multiple molecular mechanisms of muscle lipotoxicity downstream of lipid accumulation have been proposed, including oxidative stress, inflammation, insulin resistance, decreased protein synthesis, increased protein degradation, and autophagy [16]. However, the molecular mechanisms of lipid dysregulation and lipotoxicity specifically in mitochondrial myopathy, and whether they can be targeted for therapy, remain to be elucidated.

Here, we show that OxPhos defects in COX10 KO muscle trigger mitochondrial biogenesis, leading to increased, yet probably dysfunctional, pool of mitochondria with profound alterations in membrane phospholipids (PL) composition. We also show that adipose triglyceride lipase (ATGL) responsible for initiating lipolysis is upregulated in WAT and muscle of mouse models of mitochondrial myopathy and muscle of MERRF patients. Lipid metabolism remodeling in muscle results in accrual of free FA, which are inefficiently utilized in FAO and directed towards PL and ceramides (Cer) synthesis in the endoplasmic reticulum (ER). In muscle of COX10 KO mouse and muscle of human MERRF patients, we find Cer accumulation associated with chronic and progressive elevation of ER stress. We show that inhibiting *de novo* Cer synthesis decreases ER stress and ATF4-signaling, ameliorates muscle proteostasis, and improves myopathy. Together, our data indicate that in OxPhos defective muscle, chronic, unresolved induction of ER stress caused by the buildup of toxic lipid species leads to protein catabolism, thereby contributing to disease pathogenesis.

## RESULTS

### COX deficiency in muscle promotes mitochondrial biogenesis with enrichment of OxPhos complexes

In the muscle-specific COX10 KO mouse, the genetic excision of the assembly factor heme A:farnesyltransferase (COX10) occurs exclusively in skeletal muscle at embryonic stage due to activation of Myl1-Cre gene [17], resulting in a stable deficiency of COX (cytochrome oxidase, complex IV) in skeletal muscle [18]. Interestingly, although COX activity is already fully suppressed as early as postnatal day 50 and does not change further, myopathy keeps worsening over time [11, 18], similar to the progressive muscle disease observed in PMM patients with COX deficiency [19]. This suggests that, in addition to primary energy defects due to OxPhos impairment, secondary mechanisms contribute to disease progression.

Lack of heme A due to COX10 deletion impairs the assembly of functional COX complex, leading to degradation of its unassembled core subunits [20]. In accord, we found a decrease in COX subunit 1 (COXI) in COX10 KO muscle that remains stable between 100 and 200 days of age (Fig. 1A-D). At the same time, we detected increased protein levels of the UQCR2 subunit of complex III and the ATP5A subunit of complex V, at 100 and 200 days (Fig. 1A-B), and increased NDUFB8 subunit of complex I and SDHB subunit of complex II, at 200 days (Fig. 1C-D). Cytochrome *c* levels were also increased at both 100 and 200 days (Fig. S1A-B). To assess if this is the effect of compensatory OxPhos machinery biogenesis, we functionally investigated individual complexes by measuring their specific enzymatic activities. The activities of complexes I, II, and V were increased in COX10 KO muscle lysates (at 200 days, Fig. 1E-H), suggesting increased OxPhos biogenesis. Moreover, analysis of BN-PAGE followed by WB of isolated mitochondria showed an enrichment of complexes III and V at 200 days (Fig. 1I-J). Coomassie blue staining of the gels prior to transfer showed that the loading of mitochondrial protein was similar for all samples (66kDa band, Fig. S1C). To determine whether OxPhos complexes upregulation was part of a broad program of increased mitochondrial biogenesis we examined muscle mitochondrial content. Assessment of mitochondrial protein per gram of tissue revealed increased mitochondrial content (at 200 days, Fig. S1D) and was accompanied by increased levels of the mitochondrial transcription factor A (TFAM, Fig. S1E-F), which was shown to correlate with mtDNA copy number [21]. In addition, more than two-fold increase in transcript levels of peroxisome proliferator-activated receptor γ coactivator 1-α (PGC-1α), master regulator of mitochondrial biogenesis and upstream regulator of TFAM [22], were previously reported [11]. Increased mitochondrial mass in COX10 KO muscle was also evidenced by transmission electron microscopy (TEM). CTL muscle displayed a distinctive honeycomb reticular pattern with elongated intermyofibrillar mitochondria and rounder subsarcolemmal mitochondria. Instead, COX10 KO muscle showed an increased number of enlarged mitochondria with reduced density, within both the intermyofibrillar and subsarcolemmal regions (Fig. 1K-L). Ragged-red muscle fibers appear in histological staining of muscle with mitochondrial accumulation, a histopathological hallmark of human mitochondrial myopathy [23], and have been previously shown also in COX10 KO muscle [18]. Together, these findings suggest that COX10 KO muscle mitochondrial biogenesis is strongly upregulated.

**Figure 1.**
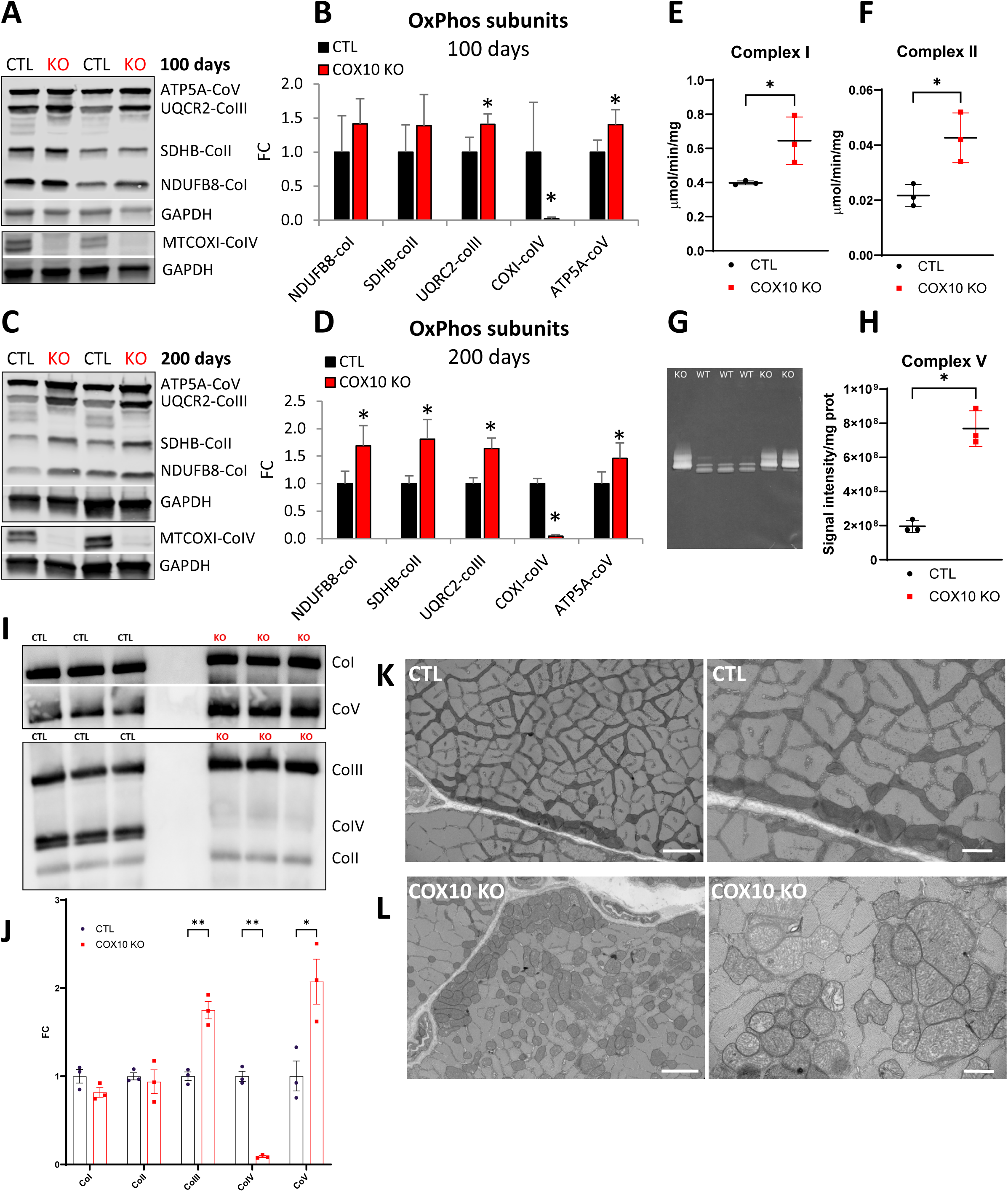
Mitochondrial biogenesis is increased in COX10 KO muscle. **A)** Representative WB of muscle lysates from 100 days old COX10 KO and CTL mice, separated by denaturing SDS-PAGE, and probed for OxPhos subunits and GAPDH. **B)** Muscle protein levels of OxPhos subunits in 100 days old COX10 KO mice (n=4), normalized by GAPDH, expressed relative to 100 days old CTL (n=4) set at 1. Data are presented as Mean ± SD, * p<0.05, by unpaired t-test. **C)** Representative WB of muscle lysates from 200 days old COX10 KO and CTL mice, separated by denaturing SDS-PAGE, and probed for OxPhos subunits and GAPDH. **D)** Muscle protein levels of OxPhos subunits in 200 days old COX10 KO mice (n=4), normalized by GAPDH, expressed relative to 200 days old CTL (n=4) set at 1. Data are presented as Mean ± SD, * p<0.05, by unpaired t-test. **E)** Complex I activity measured as NADH:HAR reductase activity in muscle homogenates from 200 days old COX10 KO (n=3) and CTL (n=3) mice, normalized by protein content. Data are presented as Mean ± SD, * p<0.05, by unpaired t-test. **F)** Complex II succinate:DCIP reductase activity measured as succinate:DCIP reductase activity in muscle homogenates from 200 days old COX10 KO (n=3) and CTL (n=3) mice, normalized by protein content. Data are presented as Mean ± SD, * p<0.05, by unpaired t-test. **G)** Representative high-resolution clear native gel after assaying complex V ATPase in gel activity in mitochondrial membranes from 200 days old COX10 KO and CTL muscle. **H)** Quantitative densitometry analysis of in-gel ATPase activity of complex V in mitochondrial membranes from COX10 KO (n=3) and CTL muscle (n=3), normalized by protein content. Data are presented as Mean ± SD, * p<0.05, by unpaired t-test. **I)** Western blot of mitochondria isolated from 200 days old COX10 KO and CTL muscle, separated by Blue Native PAGE (20 μg of mitochondrial protein /lane), and probed for NDUFA9 (complex I, CoI), SDHA (complex II, CoII), UQCRC2 (complex III, CoIII), COX1 (complex IV, CoIV), and ATPase β (complex V, CoV). **J)** Levels of OxPhos complexes in mitochondria from muscle of 200 days old COX10 KO (n=3) calculated by band densitometry, expressed relative to 200 days old CTL (n=3) set at 1. Data are presented as Mean ± SD, * p<0.05, ** p<0.01, by unpaired t-test. **K-L)** Transmission electron microscopy of gastrocnemius muscle from 200 days old CTL (K) and COX10 KO (L) mice. In (K) and (L) the images on the left panel are taken at 4000 X magnification (scale bar, 2 μm) and the images on the right panel are taken at 8000 X magnification (scale bar, 800 nm).

### COX deficiency induces a profound remodeling of muscle phospholipid profiles

Increased mitochondrial biogenesis requires the concurrent biosynthesis of new phospholipids (PL), especially the unique mitochondrial phospholipid cardiolipin (CL), which is almost exclusively localized within the inner mitochondrial membrane (IMM). CL is a dimeric phospholipid containing three glycerol backbones and four fatty acid (FA) chains and plays key roles in IMM integrity and cristae morphology [24]. CL is essential for the structure, function, and supramolecular organization of the OxPhos system [25, 26]. To investigate muscle lipidome remodeling, we utilized an integrated LC-MS and LC-MS/MS platform for comprehensive profiling. Levels of CL 72:8, a tetra linoleoyl (18:2)_4_ containing four chains of linoleic acid (LA, C18:2), an essential omega-6 PUFA, were highly increased in COX10 KO (Fig. 2A, 150 days). In addition, levels of CL species containing combinations of LA and oleic acid (OA, C18:1) such as CL 72:6 (18:1)_2_/(18:2)_2_ and CL72:7 (18:1)/(18:2)_3_, were also increased in COX10 KO muscle (Fig. 2A). Elevated levels of CL72:6, CL72:7, and CL72:8, which are the most abundant CL species in muscle mitochondria (Fig. S2A), suggest increased IMM biosynthesis upon induction of mitochondrial biogenesis. In particular, the LA-enriched CL 72:8 cardiolipin species, considered the mature form of CL, adopts a conical shape that contributes to the IMM curvature thereby supporting cristae organization and OxPhos complexes stability [27]. Therefore, the rise of CL 72:8 levels over time (Fig. 2B) likely reflects the progressive accumulation of mitochondrial mass attempting to compensate for the energetic deficit in COX10 KO muscle. On the other hand, the levels of several less abundant CL species containing docosahexaenoic acid (DHA, C24:6, omega-3 PUFA, e.g., CL 74:10, CL 74:11, CL 74:12, CL 76:10, CL 76:11, CL 76:12, CL 76:13), were decreased in COX10 KO muscle (Fig. 2C). Overall, COX10 KO muscle displayed a profound remodeling of CL profile, with increased C18:2-containing CLs and decreased C22:6-containg CLs.

**Figure 2.**
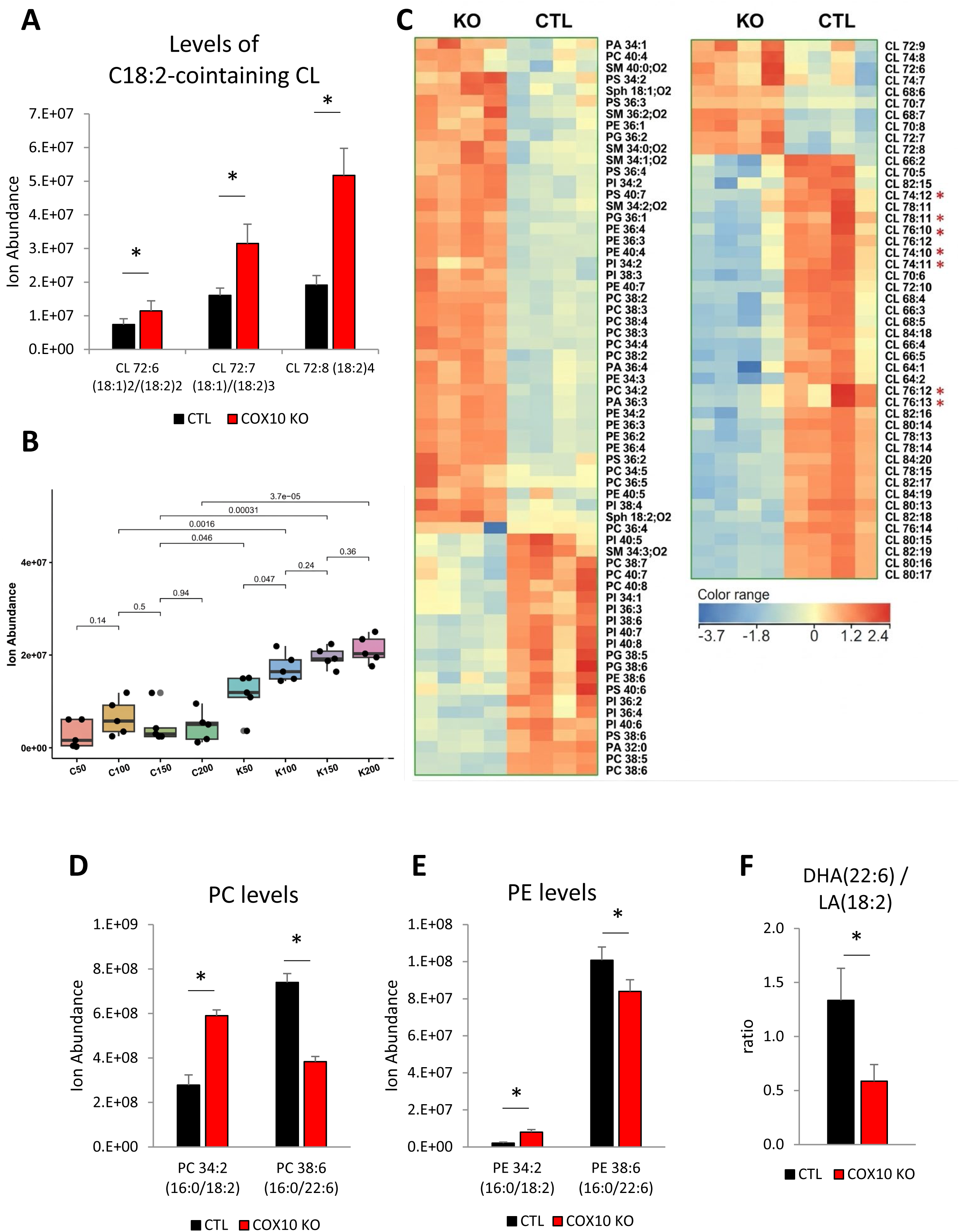
Muscle phospholipid profile is profoundly remodeled in COX10 KO mice. **A)** Muscle levels of C18:2-containing CL in 150 days old COX10 KO (n=4) and CTL (n=4) mice, measured by LC/MS analysis. Data are presented as Mean ± SD, * p<0.05, by unpaired t-test. **B)** Age dependent levels of CL 72:8 in muscle of COX10 KO (K, n=5/age group) and CTL (C, n=5/age group) mice, measured by LC/MS analysis at 50, 100, 150, and 200 days. Data are presented as Mean ± SEM. p-values, estimated by two-way ANOVA are shown for comparison among groups. **C)** Heatmap of PL and CL in COX10 KO vs. CTL muscle at 150 days, n=4 per group, by LC/MS analysis. Colors are based on the Z-score, blue denotes lower abundance and red higher abundance. Red asterisks indicate C22:6-containing CL. **D)** Muscle levels of C18:2-containing PC and C22:6-containing PC in 150 days old COX10 KO (n=4) and CTL (n=4) mice, measured by LC/MS analysis. Data are presented as Mean ± SD, * p<0.05, by unpaired t-test. **E)** Muscle levels of C18:2-containing PE and C22:6-containing PE in 150 days old COX10 KO (n=4) and CTL (n=4) mice, measured by LC/MS analysis. Data are presented as Mean ± SD, * p<0.05, by unpaired t-test. **F)** Ratio between DHA (C22:6) and LA (C18:2) in 150 days old COX10 KO (n=4) and CTL (n=4) mice, measured by LC/MS analysis. Data are presented as Mean ± SD, * p<0.05, by unpaired t-test.

Interestingly, we also detected increased levels of C18:2-containing phosphatidylcholine (PC) and phosphatidylethanolamine (PE), such as PC 34:2 and PE 34:2, and decreased levels of C22:6-containing PC and PE, such as PC 38:6 and PE 38:6 (Fig. 2D-E). In mammals, CL, PC, and PE together account for 75%–95% of mitochondrial membrane lipids [28]. Together, these results indicate that increased C18:2 incorporation is a common feature of PL remodeling in COX10 KO muscle mitochondrial membranes.

PL acyl chain composition is highly dynamic and depends on available fatty acyl chains (Acyl-CoA) and on the synthesis and remodeling of PL precursors (e.g., phosphatidic acid, PA). In the *de novo* synthesis of PL, 1-acyl-sn-glycerol-3-phosphate acyltransferase (AGPAT) converts lysophosphatidic acid (LPA) into phosphatidic acid (PA), by adding the second acyl chain (Acyl-CoA) to the *sn2*-position. AGPAT isoforms exhibit a distinct preference for specific acyl chain length and saturation, which contributes to the regulation of PL and triacylglycerols (TAG) composition, thereby influencing membrane properties [29]. AGPAT1 protein levels assessed by MS-proteomics were unchanged, and AGPAT3 and AGPAT5 upregulated in COX10 KO muscle (Fig. S2B). AGPAT1 resides exclusively in the endoplasmic reticulum (ER) and prefers short to medium saturated and monounsaturated chains (C12-C18:1), AGPAT3 in the ER and Golgi synthesizes PL containing PUFA (C18:2 and C22:6), and AGPAT5 in mitochondria has a preference for C18:1. Since we saw a decrease in C22:6 containing PL, it is unlikely that the accumulation of C18:2-containing PLs was due to AGPAT upregulation. Moreover, in the remodeling pathway (Land’s cycle), lysophospholipid (LPL) is converted into PL through addition of the acyl chain to the *sn2*-position by LPL acyltransferase (LPLAT/*Lpcat*). Protein levels of LPLAT5 (*Lpcat3)* were unchanged in COX10 KO muscle (Fig. S2C), and other LPLAT isoforms were undetected, suggesting that LPLAT is not responsible for the altered PL profile.

The increase in C18:2-containing PL and the decrease in C22:6-containing PL might reflect the relative abundance of the respective FA. DHA levels were unchanged and there was an increasing trend for LA levels (Fig. S2D) resulting in decreased DHA/LA ratio in COX10 KO muscle (Fig. 2F). Taken together, the results show profound remodeling of PL profiles in COX10 KO muscle characterized by increased levels of the most abundant CL species, which agrees with an increased need for membrane synthesis to support enhanced mitochondrial biogenesis. The molecular mechanisms behind the preferred incorporation of LA (C18:2) into PL during synthesis which primarily occurs in the ER will require further investigation.

### Increased ATGL-mediated lipid mobilization results in adipose stores depletion in COX10 KO mice

Lipids in muscle mainly derive from the uptake of circulating fats from diet and from WAT stores. The standard mouse diet contains a mixture of saturated FA [Myristic acid (14:0), Palmitic acid (16:0), and Stearic acid (18:0)], monounsaturated FA (OA, 18:1), and essential polyunsaturated FA (LA, 18:2). Dietary FA are primarily stored as TAG in WAT and released into circulation for tissue uptake and utilization as energy substrates and structural lipid biosynthesis. In both COX10 KO mice and patients affected by myoclonic epilepsy and ragged red fiber (MERRF) associated with m.8344A>G variant, we reported elevated levels of FA in plasma and muscle [11]. This suggests increased FA release into circulation from WAT and FA uptake from muscle. WAT lipolysis is initiated by the rate-limiting enzyme adipose triglyceride lipase (ATGL) which hydrolyzes stored TAG into diacylglycerol (DAG) and free FA, with cleavage at the *sn*-1 or *sn*-2 position. Consistent with increased WAT lipolysis in COX10 KO mice, we found increased levels of ATGL in both the epididymal (eWAT, visceral) and inguinal (iWAT, subcutaneous) fat depots at 200 days (40% and 92%, respectively; Fig. 3A-D). Excessive breakdown of lipid stores, not compensated by lipogenesis, results in depletion of fat mass which was detected as early as 80 days in COX10 KO mice by Echo MRI (Fig. 3E). In parallel with lower lean mass gain (Fig. 3F), fat loss contributed to failure to gain body mass at 100 days (Fig. 3G). At a later disease stage (200 days) the body mass loss further worsened (46% decrease, Fig. 3H), with an extreme loss of fat (Fig. 3I) and lean mass (Fig. 3J). A marked reduction of adipocytes size in eWAT at 200 days by H&E staining (Fig. 3K) further supported decreased fat storage capacity in COX10 KO mice. These results suggest that a progressive cachexia condition occurs alongside the primary mitochondrial dysfunction and that muscle wasting likely worsens the myopathy.

**Figure 3.**
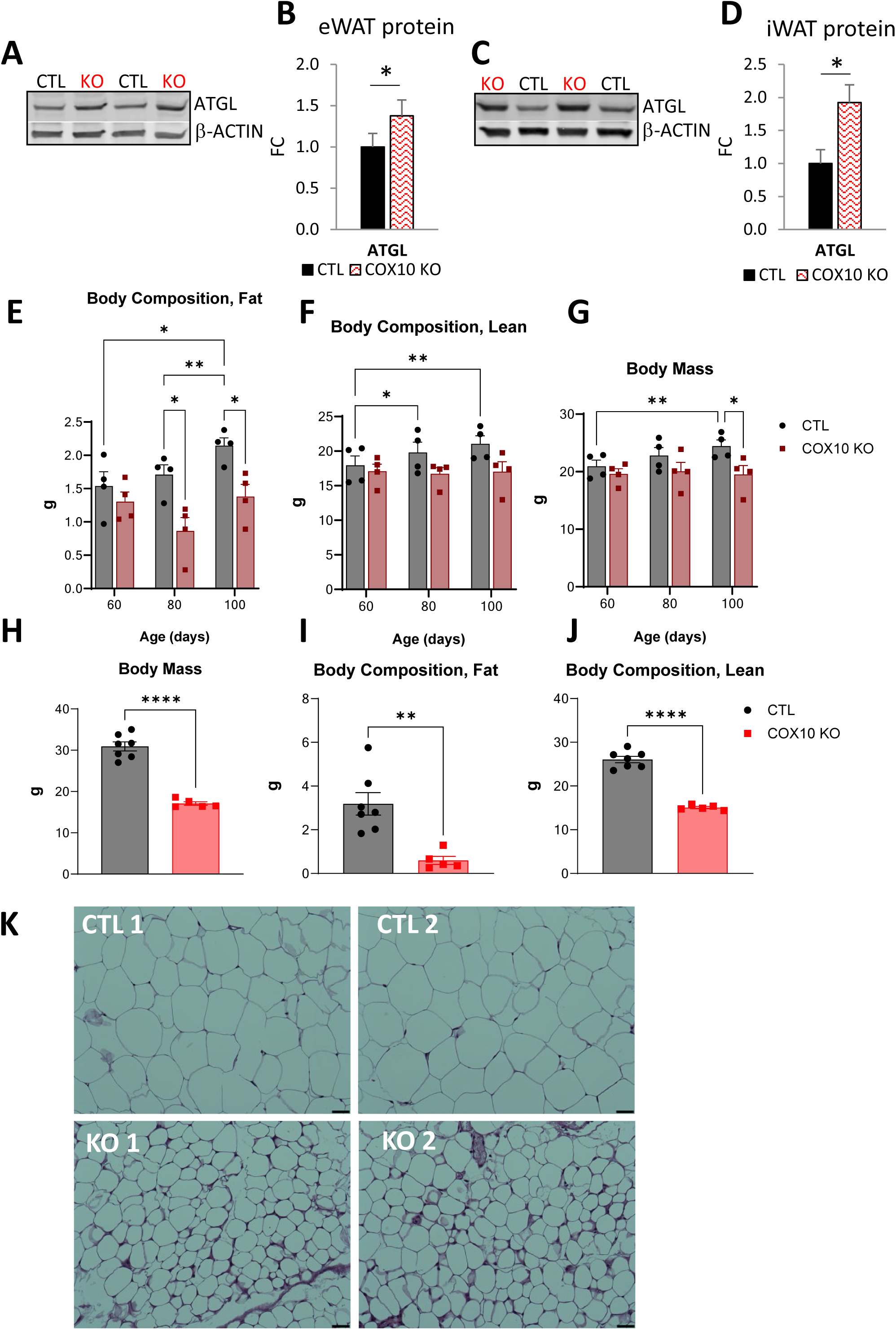
WAT stores are depleted in COX10 KO mice. **A)** Representative WB of eWAT lysates from 200 days old COX10 KO and CTL mice, separated by denaturing SDS-PAGE, and probed for ATGL and β-ACTIN. **B)** Protein levels of ATGL in eWAT of 200 days old COX10 KO mice (n=4), normalized by β-ACTIN, expressed relative to 200 days old CTL (n=4) set at 1. Data are presented as Mean ± SD, * p<0.05, by unpaired t-test. **C)** Representative WB of iWAT lysates from 200 days old COX10 KO and CTL mice, separated by denaturing SDS-PAGE, and probed for ATGL and β-ACTIN **D)** Protein levels of ATGL in iWAT of 200 days old COX10 KO mice (n=4), normalized by β-ACTIN, expressed relative to 200 days old CTL (n=4) set at 1. Data are presented as Mean ± SD, * p<0.05, by unpaired t-test. **E-G)** Age dependent Fat (E), Lean (F), and Body (G) mass in COX10 KO (n=4/age group) and CTL (n=4/age group) by Echo MRI. Data are presented as Mean ± SEM, * p<0.05, ** p<0.01 estimated by two-way ANOVA. **H-J)** Body (H), Fat (I), and Lean (J) mass in 200 days CTL (n=7) and COX10 KO (n=5) by Echo MRI. Data are presented as Mean ± SEM, ** p<0.01, **** p<0.0001 estimated by unpaired t-test. **K)** Representative image of eWAT cross sections stained with H&E from 200 days old CTL (n=2) and COX10 KO (n=2) mice. Images are taken at 20 X magnification (black scale bar, 100 μm).

### Lipids accumulate in muscle fibers in mitochondrial myopathy

Chronic fat mobilization from WAT leads to elevated circulating FA, promoting ectopic lipid accumulation in peripheral non-adipose tissues. In skeletal muscle, long-chain fatty acid (LCFA) uptake is mediated by the transmembrane transporters CD36 and FATP1 (SLC27a1), which were both increased at 200 days of age in COX10 KO muscle by proteomics analysis, relative to age-matched controls (Fig. 4A). The levels of the FA-binding proteins FABP3 (muscle isoform), FABP4, and FABP5, which are lipid chaperones facilitating LCFA transport and trafficking, were also increased in 200 days old COX10 KO muscle by proteomics (Fig. 4B). In muscle of COX10 KO mice and MERRF patients, FA accumulate in intramyocellular lipid droplets (LD) [11]. Within the LD, lipids are mainly esterified into inactive TAG, a protective mechanism that prevents free FA in the cytosol from disrupting organelle membranes [30]. However, TAG can be hydrolyzed to DAG and free FA by muscle ATGL [31], which was upregulated in COX10 KO males as early as 50 days of age (Fig. 4C-D). Interestingly, the levels of ATGL progressively increased suggesting that cytosolic FA in muscle increase with age. ATGL upregulation was also detected in 200 days old COX10 KO females (Fig. S3A-B) although earlier time points were not tested. Elevated ATGL was also found in mouse models of muscle-specific complex I (NDUFS3 KO [32]) and complex III (Rieske protein, RISP KO [33]) deficiency (Fig. 4E-H) with severe myopathy. These results indicate that, regardless of the respiratory complex affected, OxPhos impairment drives TAG hydrolysis in muscle. Importantly, we found increased levels of ATGL in MERRF muscle (Fig. 4I-J), suggesting that ATGL upregulation is a shared feature of mouse and human mitochondrial myopathy.

**Figure 4.**
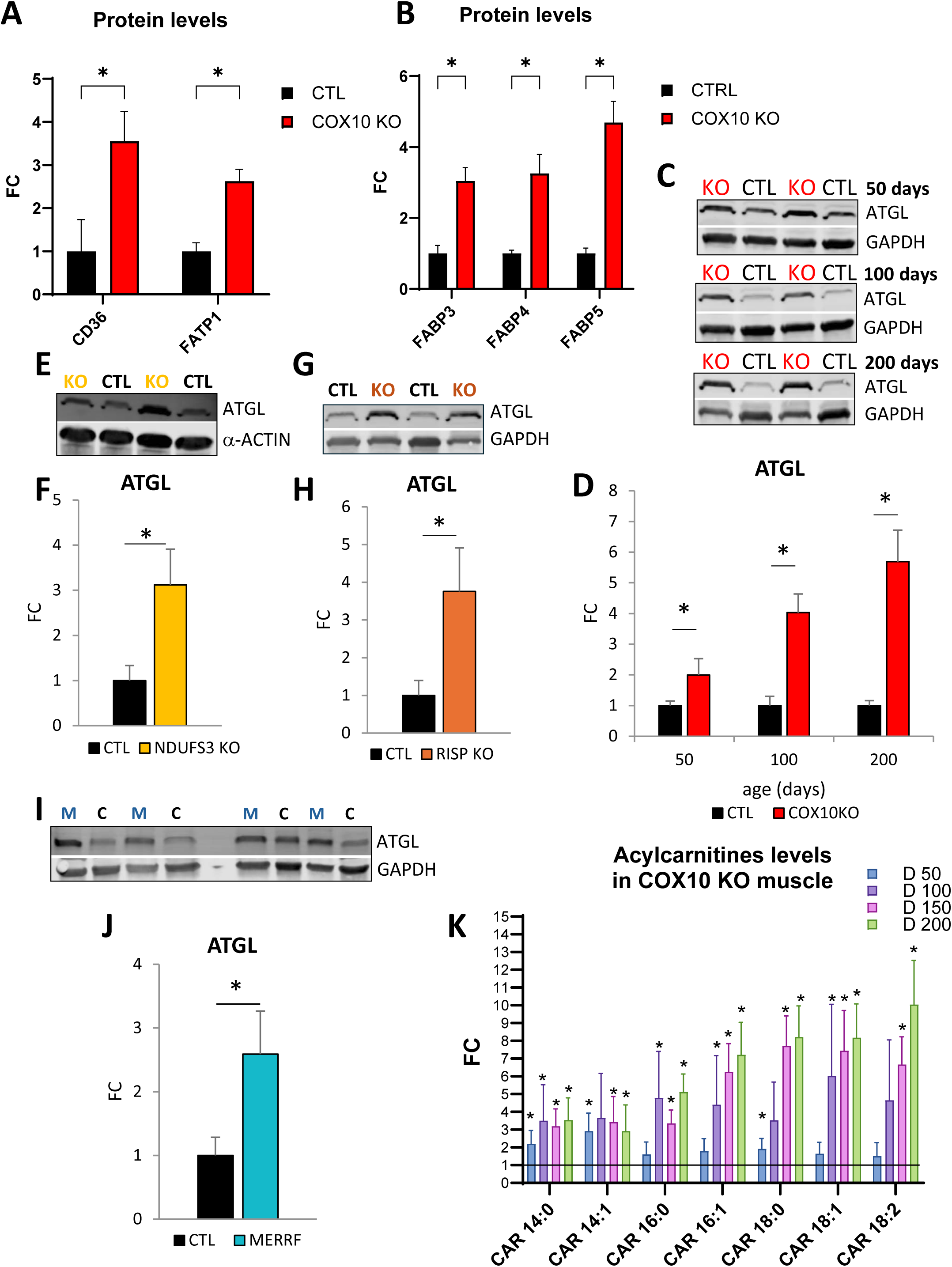
ATGL-mediated TAG breakdown is increased in OxPhos defective muscle. **A)** Muscle protein levels of CD36 and FATP1 in 200 days old COX10 KO (n=4) expressed relative to 200 days old CTL (n=4) set at 1, by proteomics analysis. Data are presented as Mean ± SD, * p<0.05, by unpaired t-test **B)** Muscle protein levels of FABP3, FABP4, and FABP5 in 200 days old COX10 KO (n=4) expressed relative to 200 days old CTL (n=4) set at 1, by proteomics analysis. Data are presented as Mean ± SD, * p<0.05, by unpaired t-test **C)** Representative WB of muscle lysates from 50, 100, and 200 days old COX10 KO and CTL mice, separated by denaturing SDS-PAGE, and probed for ATGL and GAPDH. **D)** Age-dependent muscle protein levels of ATGL in COX10 KO (n=4 per age group), normalized by GAPDH, expressed relative to CTL (n=4 per age group), at 50, 100, and 200 days. Data are presented as Mean ± SD, * p<0.05 COX10 KO vs. same age CTL set at 1, by unpaired t-test. **E)** Representative WB of muscle lysates from 100 days old NDUFS3 KO and CTL mice, separated by denaturing SDS-PAGE, and probed for ATGL and α-ACTIN. **F)** Muscle protein levels of ATGL in 100 days old NDUFS3 KO (n=4), normalized by α-ACTIN expressed relative to 100 days old CTL (n=4) set at 1. Data are presented as Mean ± SD, * p<0.05, by unpaired t-test. **G)** Representative WB of muscle lysates from 35 days old RISP KO and CTL mice, separated by denaturing SDS-PAGE, and probed for ATGL and GAPDH **H)** Muscle protein levels of ATGL in 35 days old RISP KO (n=4), normalized by GAPDH, expressed relative to 35 days old CTL (n=4) set at 1. Data are presented as Mean ± SD, * p<0.05, by unpaired t-test. **I)** WB of human muscle lysates from MERRF (M) and CTL (C), separated by denaturing SDS-PAGE, and probed for ATGL and GAPDH. **J)** Human muscle protein levels of ATGL in MERRF (n=4, 1 female and 3 males), normalized by GAPDH, expressed relative to CTL (n=4, 2 females and 2 males) set at 1. Data are presented as Mean ± SD, * p<0.05, by unpaired t-test. **K)** Muscle levels of Acylcarnitines by LC/MS analysis in COX10 KO (n=5 per age group) expressed relative to same age CTL (n=5 per age group), at 50 (D50), 100 (D100), 150 (D150), and 200 (D200) days of age. Data are presented as Mean ± SD. The black line indicates the CTL value set at 1. * p<0.05 COX10 KO vs. same age CTL by unpaired t-test

OxPhos defects cause the buildup of acyl carnitines due to the stalling of mitochondrial FA β-oxidation (FAO) associated with redox imbalance. Excessive acyl-CoA in the mitochondrial matrix is converted into acyl carnitines by mitochondrial carnitine palmitoyltransferases (CPT2), exported out of mitochondria, and eventually excreted in circulation [12, 34]. In COX10 KO muscle, we observed a progressive accumulation of acyl carnitines (C14-18 species, Fig. 4K), reflecting an overload of FA that exceeds the capacity to utilize them as energy substrates or as precursors of PL for membrane biogenesis.

### Ceramides accumulate in OxPhos defective muscle and are associated with ER stress

The accumulation of FA in skeletal muscle drives the synthesis of sphingolipid (SL) through *de novo* Ceramide (Cer) synthesis [35]. Excess Cer in muscle is linked to insulin resistance, muscular dystrophy, and sarcopenia [36–38]. Furthermore, evidence links lipotoxicity in muscle to chronic ER stress [39]. ER is the major site for the synthesis of membrane sterols, phospholipids (PL), and Cer. In the *de novo* Cer synthesis, condensation of palmitate (C16:0) and serine generates a sphingosine backbone (C18) to which a second FA of various length is attached. In very long-chain Cer (VLC-Cer), the second FA contains 22-24 carbon atoms. It was shown that high levels of VLC-Cer and DH-Cer are associated with activation of the Protein kinase R-like ER kinase (PERK)-mediated arm of the ER stress response in cultured myotubes and muscle [40]. Chronic PERK activation is implicated in decreased protein synthesis, muscle size, and insulin sensitivity [41]. Therefore, we hypothesized that in OxPhos defective muscle, FA overload could result in Cer accumulation and chronic activation of PERK-mediated ER stress, leading to muscle catabolism (Fig. 5A). In support of this hypothesis, we found that levels of muscle VLC-Cer and their precursors dihydroceramide (DH-Cer) were increased in both MERRF patients and COX10 KO mice (Fig. 5B-C). The ER-chaperone BIP (GRP78/HSPA5) is a well-established ER stress marker that is upregulated to restore protein folding homeostasis [42]. We have previously reported upregulation of BIP transcripts in COX10 KO muscle, at 200 days (1.5-fold change,[11]), and we found that BIP protein was elevated at 100 days and increased with age (Fig. 5D-E). During ER stress, the release of BIP from the luminal domain of PERK allows PERK oligomerization, autophosphorylation, and phosphorylation of the eukaryotic translation initiation factor 2 subunit alpha (eIF2α) [43]. Ph-eIF2α attenuates global protein translation while promoting the preferential translation of selected mRNAs, most notably the activating transcription factor 4 (ATF4), a basic zipper (bZIP) transcription factor for genes involved in nutrient import, metabolism, and alleviation of oxidative stress [44]. In COX10 KO muscle, we found increase in the Ph-eIF2α/eIF2α ratio (Fig. 5F-G) and ATF4 protein levels (Fig. 5H-I). The progressive accumulation of ATF4 over time (1.6-, 2.2-, and 3.7-fold change at 50, 100, and 200 days, respectively; Fig. 5H-I) could be driven by enhanced, selective translation of ATF4 and concurrent increase in ATF4 transcription [11].

**Figure 5.**
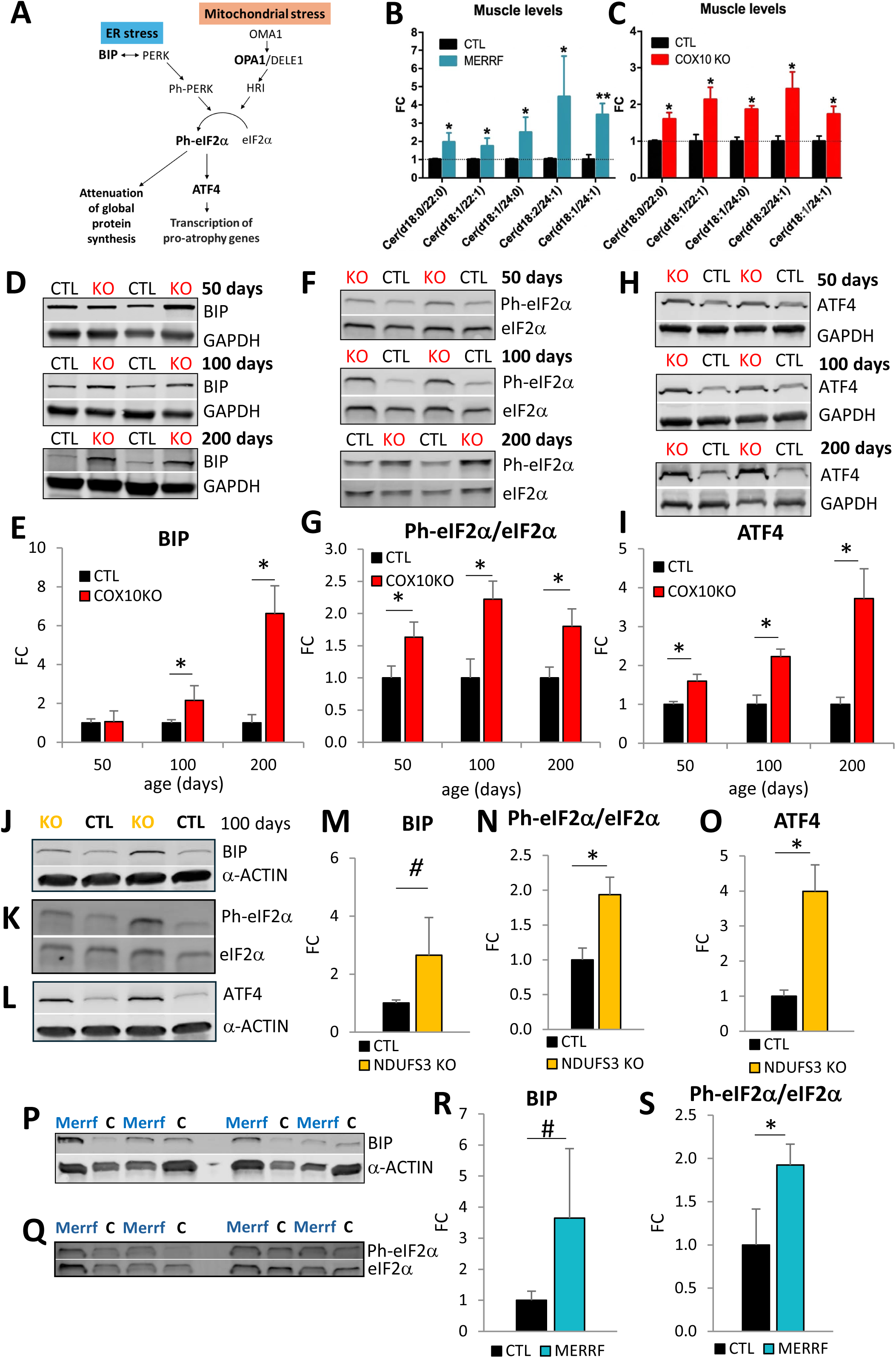
In skeletal muscle OxPhos defective mitochondria induce ER stress. **A)** Schematic representation of pathways of ER stress and mitochondrial stress converging on the ISR via phosphorylation of eIF2α. **B-C)** Quantification of a subset of muscle VLC-Cer altered in both MERRF patients (B) and COX10 KO mice (C) by LC/MS analysis. MERRF (n=4) vs. CTL (n=8). COX10 KO (n=5) vs. CTL (n=5) at 200 days. Data are presented as Mean ± SEM. The dashed line indicates CTL values set at 1. * p<0.05, ** p<0.01, by unpaired t-test. Cer (d18:1), ceramide; DHCer (d18:0), dihydroceramide. **D)** Representative WB of muscle lysates from 50, 100, and 200 days old COX10 KO and CTL mice, separated by denaturing SDS-PAGE, and probed for BIP and GAPDH. **E)** Age-dependent protein levels of BIP in COX10 KO muscle (n=4 per age group) normalized by GAPDH expressed relative to same age CTL (n=4 per age group) set at 1, at 50, 100, and 200 days. Data are as Mean ± SD, * p<0.05 COX10 KO vs. same age CTL, by unpaired t-test. **F)** Representative WB of muscle lysates from 50, 100, and 200 days old COX10 KO and CTL mice, separated by denaturing SDS-PAGE, and probed for Ph-eIF2α and eIF2α. **G)** Age-dependent levels of Ph-eIF2α/eIF2α in COX10 KO muscle (n=4 per age group) expressed relative to same age CTL (n=4 per age group) set at 1, at 50, 100, and 200 days. Data are presented as Mean ± SD, * p<0.05 COX10 KO vs. same age CTL, by unpaired t-test. **H)** Representative WB of muscle lysates from 50, 100, and 200 days old COX10 KO and CTL mice, separated by denaturing SDS-PAGE, and probed for ATF4 and GAPDH. **I)** Age-dependent protein levels of ATF4 in COX10 KO muscle (n=4 per age group) normalized by GAPDH expressed relative to same age CTL (n=4 per age group) set at 1, at 50, 100, and 200 days. Data are presented as Mean ± SD, * p<0.05 COX10 KO vs. same age CTL, by unpaired t-test. **J-L)** Representative WB of muscle lysates from 100 days old NDUFS3 KO and CTL mice, separated by denaturing SDS-PAGE, and probed for BIP and α-ACTIN (J), Ph-eIF2α and eIF2α (K), and ATF4 and α-ACTIN (L). **M-O)** Muscle protein levels of BIP normalized by α-ACTIN (M), Ph-eIF2α/eIF2α ratio (N), and ATF4 normalized by α-ACTIN (O) in NDUFS3 KO (n=4), expressed relative to CTL (n=4) set at 1, at 100 days. Data are presented as Mean ± SD, * p<0.05, # p=0.09, by unpaired t-test. **P-Q)** WB of human muscle lysates from MERRF and CTL (C), separated by denaturing SDS-PAGE, and probed for BIP and α-ACTIN (P) and Ph-eIF2α and eIF2α (Q). **R-S)** Human muscle protein levels of BIP normalized by α-ACTIN (R) and Ph-eIF2α/eIF2α ratio (S) in MERRF (n=4, 1 female and 3 males), expressed relative to CTL (n=4, 2 females and 2 males) set at 1. Data are presented as Mean ± SD, * p<0.05, # p=0.06, by unpaired t-test.

Protein levels of BIP, Ph-eIF2α/eIF2α, and ATF4 were also increased in muscle of COX10 KO female mice at 200 days of age (Fig. S3C-F). In addition, in NDUFS3 KO muscle BIP was elevated, albeit not reaching statistical significance (2.6-fold change, p=0.09, Fig. 5J and M). This was accompanied by significant increases in Ph-eIF2α/eIF2α ratio (1.9-fold change, Fig. 5K and N) and ATF4 levels (4-fold change, Fig. 5L and O). Furthermore, increased BIP (3.6-fold change, p=0.06, Fig. 5P and R), Ph-eIF2α/eIF2α (1.9-fold change, Fig. 5Q and S), and ATF4 (3-fold change, [11]) were found in MERRF human muscle, indicating that activation of the BIP-PERK-eIF2α-ATF4 pathway is common to mouse and human mitochondrial myopathy.

In addition to PERK, eIF2α can be phosphorylated by the eIF2a kinase HRI (EIF2AK1) activated by the mitochondrial integrated stress response (mtISR) through OMA1 cleavage of DELE1 [45]. Therefore, to study the putative role of HRI in the temporal activation of ATF4, we investigated OMA1-mediated cleavage of OPA1 as a marker of mitochondrial stress [46]. Unlike BIP, which increases over time, OPA1 cleavage does not change with age and disease progression in COX10 KO muscle (Fig. 6A-B). This suggests that in COX10 KO muscle mtISR is initiated early on and remains stable over time, while ER stress builds up as the disease advances.

**Figure 6.**
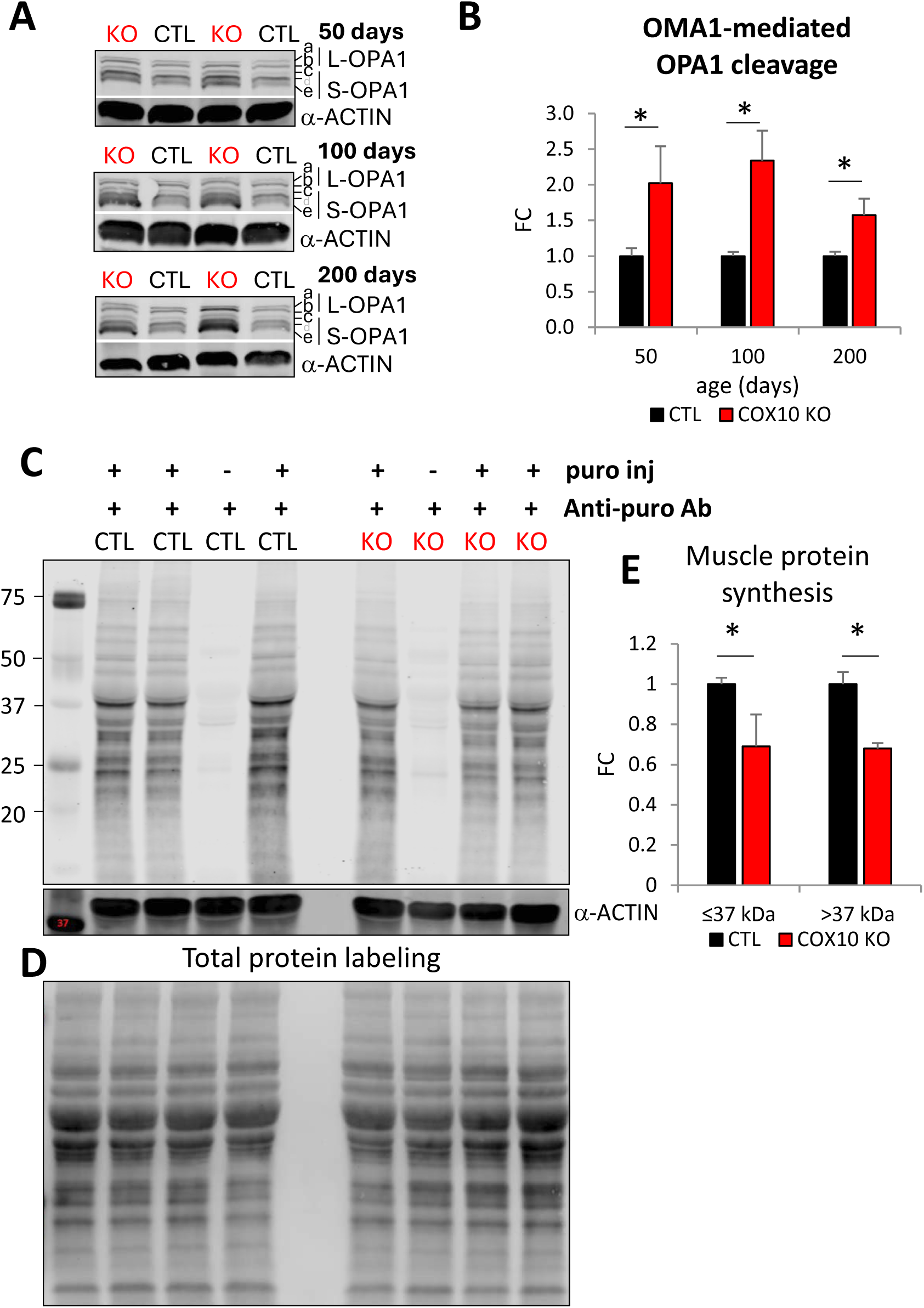
Global protein synthesis is decreased in COX10 KO muscle. **A)** Representative WB of muscle lysates from 50, 100, and 200 days old COX10 KO and CTL mice, separated by denaturing SDS-PAGE, and probed for OPA1, SDHA, CS, and α-ACTIN. **B)** Age-dependent protein levels of OMA1-mediated OPA1 cleavage calculated as S-OPA1/L-OPA1= (c+e) / (a+b)[46]) in COX10 KO muscle (n=4 per age group) expressed relative to same age CTL (n=4 per age group) set at 1, at 50, 100, and 200 days. Data are presented as Mean ± SD, * p<0.05 COX10 KO vs. same age CTL, by unpaired t-test. **C)** WB of muscle lysates from 100 days old CTL and COX10 KO mice injected (+) or not (-) with puromycin, separated by denaturing SDS-PAGE, and probed for anti-puromycin and α-ACTIN. **D)** Total proteins detected by No-StainTM protein labeling reagent in the membrane shown in C. **E)** Levels of newly synthesized muscle proteins in COX10 KO (n=3), normalized by total protein, expressed relative to CTL (n=3) set at 1. Data are presented as Mean ± SD, * p<0.05, by unpaired t-test.

Phosphorylation of eIF2α blocks the exchange of eIF2-GDP to eIF2-GTP, thus reducing global translation initiation and protein synthesis [43]. To investigate protein synthesis in muscle we performed *in vivo* Surface sensing of translation (SUnSET) assay, which allows the assessment of global protein synthesis rates through puromycin incorporation into nascent polypeptides [47]. Briefly, 100 days old mice were injected with puromycin (0.04 μmol/g in 100 μl of PBS) or PBS and then sacrificed after 30 minutes to evaluate the incorporation of puromycin in nascent muscle proteins by WB with anti-puromycin antibodies. We found that muscle protein synthesis was attenuated in COX10 KO (Fig. 6C-E). Together, these results suggest that ER stress inhibits global protein synthesis which could contribute to muscle wasting in COX10 KO.

### Inhibition of Cer synthesis decreases ER stress and improves mitochondrial myopathy

To study the causal relationship between increased Cer levels and ER-stress COX10 KO mice were treated with myriocin, an irreversible high-affinity inhibitor of Serine palmitoyltransferase (SPT), the first and rate-limiting step of *de novo* Cer synthesis. Myriocin (0.4 mg/Kg, on alternate days, [38]) or vehicle (PBS with 0.4% BSA) were injected intraperitoneally (IP) in 120 days old COX10 KO mice for 10 days. CTL mice received vehicle only. As expected, myriocin decreased the levels of several Cer precursors (DH-Cer) and VLC-Cer in muscle of COX10 KO mice compared to untreated COX10 KO (Fig. S4A). In addition, myriocin treatment also decreased BIP (Fig. S4B-C), supporting the hypothesis that Cer accumulation contributes to ER-stress in COX10 KO muscle.

Next, to investigate the phenotypic effect of Cer synthesis inhibition we treated COX10 KO mice with myriocin (0.4 mg/Kg, in PBS with 0.4% BSA, on alternate days, by IP injection) for 3 months starting at 60 days of age. By the end of the treatment, COX10 KO mice that received myriocin showed increased BW (17%, Fig. 7A), gastrocnemius mass (44%, Fig. 7B), quadricep mass (27%, Fig. 7B), and iWAT mass (52%, Fig. 7C), compared to vehicle treated COX10 KO mice. Importantly, myriocin improved COX10 KO grip strength (21%, Fig. 7D) and endurance exercise by treadmill (73%, Fig. 7E). Myriocin did not affect BW, muscle strength, and endurance in CTL mice; only gastrocnemius mass was slightly increased (17%, Fig. 7B). We confirmed that myriocin decreased DH-Cer and Cer levels in COX10 KO muscle (Fig. 7F). Myriocin did not affect Cer levels in CTL muscle, although some Cer-derived sphingomyelin and glycosphingolipid species (i.e., SM 36:1, SM 32:2, and GM3 36:1) were decreased (Fig. S4D). Furthermore, there was a trend towards reduced BIP (25%) and significantly decreased ATF4 (20%) in myriocin treated COX10 KO muscle (Fig. 7G-H), further suggesting that attenuation of Cer synthesis reduces ER stress in COX10 KO muscle.

**Figure 7.**
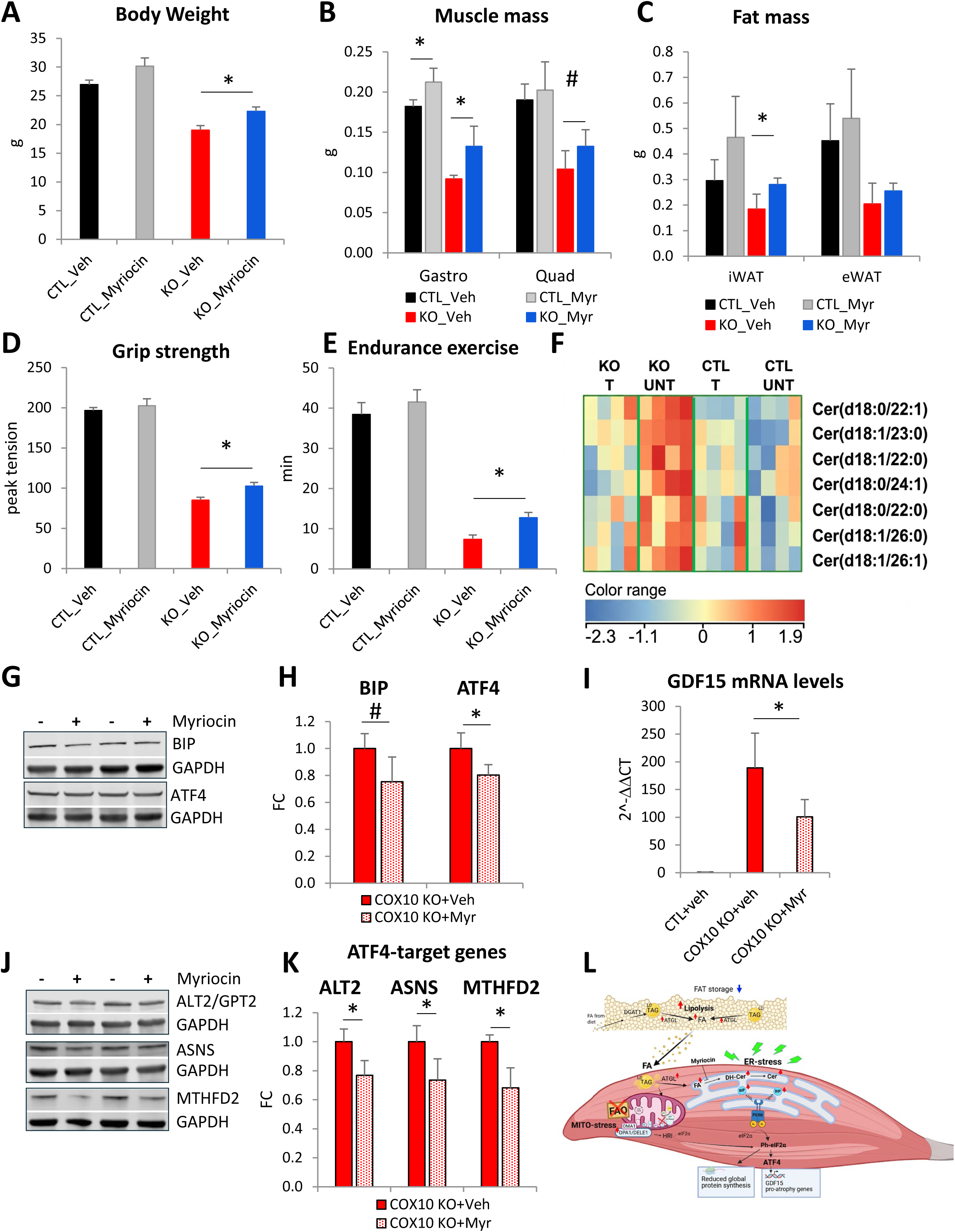
Inhibiting *de novo* Cer synthesis decreases ER stress and improves myopathy. **A-E)** BW (A), Muscle mass (B), WAT mass (C), Grip strength (D), and Endurance exercise (E) in 150 days old COX10 KO and CTL mice treated for 3 months with myriocin (n=4/genotype) or vehicle (n=5/genotype). Data are presented as Mean ± SD, * p<0.05, # p <0.1, by unpaired t-test. **F)** Heatmap of muscle Ceramide (d18:1) and dihydroceramide (d18:0) after 3 months of myriocin treatment (T) in COX10 KO and CTL compared to COX10 KO and CTL mice treated with vehicle (UNT), at 150 days, by LC/MS analysis. n=4 per treatment/per genotype. Colors are based on the Z-score, blue denotes lower abundance and red higher abundance. **G)** Representative WB of muscle lysates from COX10 KO mice treated with myriocin or vehicle, separated by denaturing SDS-PAGE and probed for BIP, ATF4, and GAPDH. **H)** Muscle protein levels of BIP and ATF4 normalized by GAPDH in COX10 KO treated with myriocin (n=4) expressed relative to COX10 KO treated with vehicle (n=4) set at 1. Data are presented as Mean ± SD, * p<0.05, # p=0.06, by unpaired t-test. **I)** GDF15 mRNA levels in muscle of COX10 KO treated with myriocin (n=4) and vehicle (n=4) and CTL with vehicle (n=3) at 150 days, by qRT-PCR, calculated by 2^-ΔΔCt^, with β-actin normalization and relative to CTL value set at 1. Data are presented as Mean ± SD, * p<0.05, by unpaired t-tests. **J)** Representative WB of muscle lysates from COX10 KO mice treated with myriocin or vehicle, separated by denaturing SDS-PAGE, and probed for ATL2, ASNS, MTHFD2, and GAPDH. **K)** Muscle protein levels of ATL2, ASNS, and MTHFD2 normalized by GAPDH in COX10 KO with myriocin (n=4) expressed relative to COX10 KO vehicle (n=4) set at 1. Data are presented as Mean ± SD, * p<0.05, by unpaired t-test. **L)** Schematics of muscle and WAT metabolic remodeling contributing to the pathogenesis of mitochondrial myopathy. Blue arrows indicate down- and red arrow up- regulation.

Chronic, unresolved activation of ATF4 leads to muscle atrophy [48, 49]. Among the pro-atrophy ATF4-target genes is the myokine growth differentiation factor 15 (GDF15) which suppress food intake, promotes WAT lipolysis, and increases muscle energy expenditure [50, 51]. We have previously shown that upregulation of GDF15 transcription in COX10 KO muscle results in progressive increase in plasma GDF15 levels [11]. Moreover, plasma GDF15 levels in PMM patients correlate with disease severity [52]. Interestingly, myriocin reduced GDF15 transcript levels in muscle of COX10 KO mice (47%, Fig. 7I), suggesting that ATF4 signaling can be modulated by Cer synthesis inhibition. In accord, additional ATF4 targets, such as ALT2, ASNS, and MTHFD2 were decreased at the protein level (23%, 26%, and 32%, respectively, Fig. 7H-I). Together, these results suggest that Cer synthesis inhibition ameliorates myopathy by attenuating ER stress and ATF4 signaling.

## DISCUSSION

Understanding tissue-specific adaptation to mitochondrial defects is essential for developing therapies for mitochondrial diseases. In this study, we utilized mouse models of mitochondrial myopathy with OxPhos defects restricted to skeletal muscle to distinguish muscle-autonomous metabolic rewiring from systemic effects. These models show that a multi-stage metabolic remodeling initiated by primary muscle mitochondrial dysfunction evolves into a systemic catabolic state resulting in progressive muscle degeneration. Compensatory mitochondrial biogenesis triggered by OxPhos defects is a common response in muscle of PMM patients and mouse models of mitochondrial myopathy. Although mitochondrial biogenesis is believed to be adaptive, direct evidence of its clinical benefit in mitochondrial myopathy is lacking [53]. In OxPhos defective muscle, activation of energy (AMPK) or redox sensing (Sirtuins) pathways upregulate PGC-1α, a master regulator of mitochondrial biogenesis, in the attempt to boost residual OxPhos capacity [54, 55]. Beyond the transcriptional upregulation of mitochondrial and nuclear genes, mitochondrial biogenesis also requires increased FA to provide the acyl groups that support mitochondrial membrane biosynthesis. Thus, the upregulation of FA availability in muscle could be crucial for mitochondrial biogenesis. It was recently shown that in *S. cerevisiae* and mammalian cells TAG lipases are essential to recover from mitochondrial stress by releasing acyl groups for CL biosynthesis needed to replenish mitochondrial content [56]. Here, we show that ATGL is upregulated in COX10 KO, NDUFS3 KO, RISP KO, and human MERRF muscle (Fig. 4) suggesting that, regardless of the complex affected, upregulation of muscle ATGL could support mitochondrial biogenesis, even when FAO capacity is limited. Therefore, increased ATGL could represent a mitochondrial biogenesis signature of OxPhos defective muscle shared by humans and mice. Future studies inhibiting muscle ATGL could directly test its role in mitochondrial biogenesis.

In the mitochondrial biogenesis process in COX10 KO muscle complexes III and V are significantly upregulated (Fig. 1). Complex III upregulation coupled with increased levels of cytochrome *c* (Fig S1A-B) could be involved in attempting to maintain electron transport when complex IV is depleted, which would require alternative pathways for cytochrome *c* oxidation. It was previously shown that during anoxia complex IV can be bypassed and cytochrome *c* oxidized by alternative redox enzymes [57]. A potential candidate is the redox protein p66Shc which is localized within the mitochondrial intermembrane space and mediates electron transfer and cytochrome *c* oxidation [58]. Moreover, during hypoxia reduction of fumarate can sustain the input of electrons by complex I thus enabling NADH reoxidation [59]. Whether alternative redox reactions are also involved in COX10 KO muscle remains to be investigated.

During fasting, the skeletal muscle PL profile is characterized by decreased DHA (22:6)-containing PC and increased LA (18:2)-containing PC [60], which is similar to the lipid profile of COX10 KO muscle (Fig. 2). It was shown that pharmacological induction of mitochondria proliferation results in decreased hepatic DHA due to increased peroxisomal DHA-CoA oxidase [61]. In COX10 KO muscle, levels of DHA are unchanged despite the DHA/LA ratio decrease, suggesting that increased DHA degradation is unlikely in this lipid remodeling. In sarcopenic muscle, upregulation of the FA chaperone FABP3 has been linked to the decrease of polyunsaturated PL, such as DHA-containing PL, which confers membrane rigidity [62]. Moreover, this FABP3-induced lipid remodeling activates PERK-mediated ER stress and inhibits protein synthesis in sarcopenic muscle. Interestingly, we find elevated levels of FABP3, FABP4, and FABP5 in COX10 KO muscle (Fig. 4), suggesting that selective affinity for certain FA by FABPs could participate in PL remodeling and alterations of mitochondrial membrane morphology.

In skeletal muscle, when FA from circulation or lipid droplets exceed the metabolic capacity for oxidation and storage, they are directed toward alternative pathways that could participate in lipotoxicity. In particular, accumulation of palmitic acid and its active form palmitoyl-CoA drive *de novo* Cer synthesis in the ER through a four-step process [63]. The first and rate-limiting step is the condensation of L-serine and palmitoyl-CoA by serine palmitoyltransferase (SPT), followed by the reduction to sphinganine, N-acylation by ceramide synthases (CerS1-6) to form dihydroceramide, and desaturation to Cer. In mammals, the length of the second Cer acyl chain is determined by the specific ceramide synthase isoform [64]. In both MERRF patients and COX10 KO muscle, VLC-Cer (C20-26) are increased (Fig. 5), suggesting that they are synthesized by CerS2. The concomitant elevation of DH-Cer further suggests that Cer are being produced *de novo* in COX10 KO muscle, and not through sphingomyelinase (SMase) or the salvage pathways or imported from circulation.

It was shown that high levels of VLC-Cer and DH-Cer are associated with activation of PERK signaling in cultured myotubes and muscle [40]. Moreover, Cer and DH-Cer can activate the ER stress sensors via transmembrane domains that are independent from the luminal domain responsible for proteotoxic stress activation [65, 66]. Therefore, perturbations of ER membrane lipid composition can activate ER stress through mechanisms independent of misfolded proteins [67]. Here, we show that in COX10 KO mice SPT inhibition with myriocin decreases ER stress and ATF4 signaling and improves muscle function. Importantly, myriocin increases both muscle and WAT mass. Our results support the hypothesis that in OxPhos defective muscle excess production of Cer drives maladaptive ER stress, contributing to the progression of mitochondrial myopathy. A potential limitation of SPT inhibition is that it can lead to the buildup of free PA or palmitoyl-CoA, which could be shunted towards other damaging lipid species, such as *sn1,2* DAG, saturated FA, LC Acyl CoA, and lysophosphatidylcholine, which have been shown to induce ER stress [68]. Therefore, inhibiting the downstream PERK signaling pathway could be an alternative strategy for mitigating muscle lipotoxicity.

In *C. elegans,* proteotoxic stress is managed by a coordination between mitochondria and ER stress with induction of the PERK-eIF2α signaling, which reduces protein load by suppressing global protein translation [69]. Interestingly, during proteotoxic stress the signaling axis from mitochondria to ER is mediated by alteration of lipid metabolism, in particular perturbations of Cer and CL synthesis [70]. On the other hand, in *Drosophila* models of Parkinson’s disease, activation of ER stress by defective mitochondria is neurotoxic [71]. The suppression of PERK signaling prevents neurodegeneration, suggesting activation of ER stress, when chronic, may contribute to neuronal dysfunction, rather than protection.

Here, for the first time in mammalian muscle, we demonstrate that OxPhos defective mitochondria induce ER stress through a lipid-mediated mechanism. Evidence shows that FAO inhibition in muscle (muscle-specific Cpt1b KO mouse) leads to accumulation of lipid droplets, lipotoxic Cer, and increased mitochondrial biogenesis [72]. Therefore, we speculate that a mitochondrial-ER signaling axis could involve abnormal accumulation of intracellular lipids. The ER increases PL synthesis to support mitochondrial biogenesis, enhance FAO, and restore lipid homeostasis. However, in OxPhos defective muscle, this adaptive mechanism fails, because incomplete FAO leads to a buildup of lipotoxic intermediates, which cause chronic ER stress, and further exacerbate myopathy. How the ER senses lipid alterations needs to be elucidated but could involve both the composition of the ER lipid bilayer and the ER-mitochondria contact sites [73].

In the COX10 KO mouse, a progressive decrease of fat mass occurs in parallel to muscle wasting (Fig. 3). Increased ATGL in both iWAT and eWAT along with adipocytes atrophy strongly suggest that adipose stores depletion is primarily driven by enhanced lipolysis rather than defective adipogenesis (adipocyte development) or hypoplasia (reduced adipocyte number). Adipose tissue is not directly affected by COX10 deletion, suggesting that a systemic signaling initiated by OxPhos defective muscle drives the mobilization of lipids from WAT. Therefore, WAT lipolysis could contribute to muscle lipotoxicity through a FA spillover mechanism that contributes to muscle FA overload. We have previously shown that in COX10 KO mice WAT wasting is mediated by the leptin-glucocorticoid signaling [11]. Moreover, it was recently shown that inhibiting GDF15 signaling in mitochondrial myopathy (mtDNA mutator, POLG mouse) ameliorates muscle function through reduction of glucocorticoid signaling [74]. This evidence suggests that modulation of GDF15-glucocorticoid signaling could be a viable strategy to attenuate the lipotoxic component of mitochondrial myopathy.

## MATERIALS AND METHODS

### Patients

Skeletal muscle biopsies from MERRF patients and controls were collected and stored in the Fondazione Policlinico Universitario A. Gemelli IRCCS, Rome, Italy. Informed consent was obtained from each participant, and the study was approved by the Ethics Committee of the Università Cattolica del Sacro Cuore (Rome, Italy; ID: 3754). Biopsy of deltoid muscle collection were performed after eight hours of fasting, for diagnostic purpose, at symptomatic stage, at the Fondazione Policlinico Universitario A. Gemelli IRCCS. Patients did not follow special diets. All patients underwent extensive clinical and radiological evaluations. Patients and controls information can be found in Table S1.

### Animals

All animal procedures were conducted in accordance with Weill Cornell Medicine Animal Care and Use Committee and performed according to the Guidelines for the Care and Use of Laboratory Animals of the National Institutes of Health. Homozygous COX10^flx^ mice, a generous gift from Dr. Francisca Diaz (University of Miami), were crossed with muscle-Cre mice (*Myl1^tm1(cre)Sjb^*/J, RRID:IMSR_JAX:024713) to generate the skeletal muscle specific COX10 KO mice. The phenotype of these mice has been described previously [18]. Only male mice were utilized for our experiments and data analysis, unless otherwise indicated. Mice were euthanized by cervical dislocation for bioenergetic/biochemical assays and with carbon dioxide for proteomics and lipidomics studies. For immunohistochemistry and transmission electron microscopy, mice were euthanized with sodium pentobarbital (150 mg/kg, i.p.). Gastrocnemius muscle, quadriceps muscle, epididymal white adipose tissue (eWAT), and inguinal white adipose tissue (iWAT) were collected from all animals at the same time of the day (10 am-11 am). All muscle experiments and data analysis were performed on gastrocnemius, unless otherwise indicated.

### Myriocin treatment

COX10 KO and CTL mice were treated with myriocin (Sigma M1177, 0.4 mg/Kg, by IP, on alternate days) or vehicle (PBS with 0.4% BSA) for 3 months starting at 60 days of age. Mice were evaluated for body weight, grip strength, and exercise endurance (by treadmill). Tissues collected at the end of the treatment (150 days of age) were weighed, frozen in liquid nitrogen, and stored at -80°C.

### Endurance Exercise

Endurance exercise was measured as time to fatigue on a conventional treadmill running task. The treadmill was equipped with a motivational grid (Columbus Instruments). After three training sessions, mice were placed on the treadmill at a fixed 5% incline with a speed starting at 5 m/min increasing incrementally by 1 m every 3 min up to 11 m/min and then and by 1 m every 1 min, as previously described [75]. Time of fatigue was recorded once mice failed to maintain pace.

### Muscle strength

Forepaw muscle strength was measured using a digital grip-strength meter (Columbus Instruments). Animals were trained to grasp a horizontal grasping ring. The testing was repeated 3 consecutive times at 15 min intervals and the average was recorded [76].

### Body composition

The total body composition of mice was measured using an EchoMRI 3-in-1 Body Composition Analyzer (EchoMRI LLC, Houston, TX) at the Metabolic Phenotyping Center, Weill Cornell Medicine (WCM). Conscious mice were placed individually into a clear plastic holding tube and analyzed without anesthesia. Total body fat mass, lean mass, free water, and total body water were quantified under 3 minutes/mouse by quantitative magnetic resonance according to the manufacturer’s instructions. Measurements were performed at the indicated experimental time points, and mice were returned to their home cages immediately after analysis.

### Immunohistochemistry

Mice were terminally anesthetized with sodium pentobarbital (150 mg/kg, i.p.) and perfused intracardially with phosphate-buffered saline (PBS). WAT depots were post-fixed in 10% neutral-buffered paraformaldehyde (PFA, Sigma) overnight at 4 °C followed by 70% ethanol before being paraffin embedded. WAT was sectioned at 5 μm on a Rotary Microtome. Paraffin-embedded tissue sections were deparaffinized by incubating the slides in Histo-Clear twice for 5 minutes each. The sections were then rehydrated through a graded ethanol series consisting of 100% ethanol twice for 3 minutes each, followed by 95% ethanol for 3 minutes and 75% ethanol for 1 minute. Following deparaffinization and rehydration, hematoxylin and eosin (H&E) staining was performed using standard histological procedures as previously described [77].

### Transmission electron microscopy

Mice were terminally anesthetized with sodium pentobarbital (150 mg/kg, i.p.) and perfused intracardially with PBS. Next, mice were perfused with 4% PFA. Gastrocnemius muscle was dissected and immersed in fixative solution (4% PFA, 2.5% glutaraldehyde in 0.1 M sodium-cacodylate). Processing, embedding, and sectioning for EM imaging was performed by the Microscopy and Image Analysis Core at WCM. Images were taken on the Hitachi HT7800 Electron microscope at the WCM Neuroanatomy Electron Microscopy Core.

### Protein analysis

#### SDS-PAGE

Skeletal muscle and WAT were rapidly dissected, snap frozen in liquid nitrogen, and kept at -80°C for western blot analysis. Tissues (approximately 20 mg) were pulverized in liquid nitrogen, with porcelain mortar and pestle (CoorsTek) placed on dry ice and suspended in RIPA buffer (500 μl) supplemented with protease inhibitors (Sigma-Aldrich, 11697498001) and phosphatase inhibitors (100x, NaF 100 mM, Na_3_VO_4_ 100 mM, Pyrophosphate 100 mM, Imidazole 200 mM, pH 7). After 30 min incubation on ice, tissue homogenate was centrifuged at 15,000 x g for 20 min at 4°C and supernatant collected. Protein concentration was determined using the DC Protein Assay (BioRad, #5000112). For electrophoresis, proteins (50 μg) were suspended in Laemmli buffer (BioRad, #1610737) containing β-mercaptoethanol, loaded on Any kD Mini-Protean TGX protein gels (BioRad, #456-8124), and separated by SDS-PAGE. Proteins were transferred to PVDF membranes (BioRad, #1704275) using a Trans-Blot® Turbo-transfer system (BioRad) and blocked in Intercept Blocking Buffer (LI-COR Biosciences, #927-60001). Membranes were probed overnight at 4°C with specific primary antibodies against the proteins of interest: anti-total OXPHOS Rodent WB Antibody Cocktail (Abcam Cat# ab110413, RRID:AB_2629281), anti-NDUFA9 CoI (Thermo Fisher Scientific Cat# 459100, RRID:AB_10376187), anti-SDHA CoII (Thermo Fisher Scientific Cat# 459200, RRID:AB_10838019), anti-UQCRC2 CoIII (Abcam Cat# ab14745, RRID:AB_2213640), anti-MTCO1 CoIV (Thermo Fisher Scientific Cat# 459600, RRID:AB_10374492), anti-β subunit CoV (Thermo Fisher Scientific Cat# A-21351, RRID:AB_221512), anti-TFAM (Santa Cruz Biotechnology Cat# sc-166965, RRID:AB_10610743), anti-ATGL (Proteintech Cat# 55190-1-AP, RRID:AB_11182818), anti-BIP (BD Biosciences Cat# 610978, RRID:AB_398291), anti-eIF2α (Cell Signaling Technology Cat# 9722, RRID:AB_2230924), anti-Ph-eIF2α (Abcam Cat# ab32157, RRID:AB_732117), anti-ATF4 (Proteintech Cat# 60035-1-Ig, RRID:AB_2058598), anti-Cytochrome *c* (BD Biosciences Cat# 556433, RRID:AB_396417), anti-GAPDH (Proteintech Cat# 60004-1-Ig, RRID:AB_2107436), anti-αACTIN (Abcam Cat# ab88226, RRID:AB_11143753), anti-βACTIN (Proteintech Cat# 66009-1-Ig, RRID:AB_2687938), anti-αTUBULIN (Proteintech Cat# 66031-1-Ig, RRID:AB_11042766), anti-TIM23 (BD Biosciences Cat# 611222, RRID:AB_398754), anti-ASNS (Proteintech Cat# 14681-1-AP, RRID:AB_2060119), anti-MTHFD2 (Proteintech Cat# 12270-1-AP, RRID:AB_2147525), anti-ALT2/GPT2 (Proteintech Cat# 16757-1-AP, RRID:AB_2112098), anti-puromycin (Sigma-Aldrich Cat# MABE343, RRID:AB_2566826). For protein detection, membranes were incubated for 1 hour at room temperature with IRDye 800CW donkey anti-mouse (LI-COR Biosciences, #926-32212) or IRDye 680RD donkey anti-rabbit (LI-COR Biosciences, # 926-68073) IgG secondary fluorescent antibodies (1:15000 dilution) and imaged using the LI-COR CLx imaging system. Band intensities were quantified using Image Studio software (vs3.1, LI-COR Biosciences). GAPDH, α-ACTIN, β-ACTIN, and α-TUBULIN were used to normalize proteins of muscle and liver homogenates.

#### Blue Native PAGE

Muscle mitochondria were isolated and solubilized as previously described [11]. Native proteins (20 μg mitochondrial protein/lane) were separated on 4-16% NativePage gradient gel (Invitrogen). Electrophoresis was performed as described [78]. After overnight transfer of native protein onto PVDF membrane at 30V, membrane was incubated overnight at 4°C with monoclonal antibodies anti-NDUFA9 CoI, anti-SDHA CoII, anti-UQCRC2 CoIII, anti-MTCO1 CoIV, anti-β subunit CoV. For immunodetection of mitochondrial complexes membrane was incubated for 1 hour with anti-mouse HRP-conjugated secondary antibodies (Jackson Immunoresearch, #115-035-146) and imaged by chemiluminescence on a ChemiDoc system (BioRad).

#### IV-SUnSET assay

*In vivo* Surface SEnsing of Translation (IV-SUnSET) technique was used to measure global protein synthesis as described [47]. Briefly, mice were i.p. injected with puromycin 0.04 μmol/g of BW dissolved in 100 μl of PBS. After exactly 30 min, muscle tissue was collected, frozen, and then processed for WB using mouse monoclonal anti-puromycin antibody (clone 12D10). The reactivity to puromycin incorporation into newly synthesized proteins was used as a measure of global protein synthesis.

### LC/MS lipidomics

LC-MS grade acetonitrile (ACN), isopropanol (IPA) and methanol (MeOH), 1-butanol (BuOH) were purchased from Fisher Scientific. High purity deionized water was filtered from Millipore (18 OMG). Lipid standards were purchased from Avanti Polar Lipids (Alabaster, AL, USA). Ammonium formate was obtained from Sigma-Aldrich in the best available grade.

Tissue lipids were extracted by adding 500µl 50:50 MeOH:BuOH + 10mM NH4Formate. The tissue–solvent mixture was subjected to bead-beating for 45 sec using a Tissuelyser cell disrupter (Qiagen), followed by 15-min sonication at 20°C. The mixture was centrifuged for 10 min at the maximum speed. Repeat the above procedures two more times to pool the supernatant. The final lipid extracts were dried in a Vacufuge (Eppendorf) and stored at -80°C until analysis. The protein pellet was solubilized in 0.2 M NaOH at 95°C for 20 min and protein was quantified using a BioRad DC assay. On the day lipidomics sample run, dry-down lipids were reconstituted in 50:50 MeOH:BuOH + 10mM NH4 Formate by normalizing to tissue protein content at 2µg/ul protein. vortexed and sonicated to solubilize the lipids. The supernatant was transferred to LC sample vials for positive and negative ion LC/MS lipidomics data acquisition. Separately, pooled quality control samples were prepared by mixing an aliquot (5ul) of each reconstituted sample extract to be used for LC MS/MS data acquisition in positive and negative ion modes.

The LC/MS platform for lipidomics analysis was described previously (Dumelie et al, 2023, PMID: 37973889), consisting of an Agilent Model 1290 Infinity II liquid chromatography system coupled to an Agilent 6550 iFunnel time-of-flight MS analyzer. An Agilent ZORBAX Eclipse Plus C18, 100 × 2.1 mm, 1.8 µm (Cat# 959758-902) reversed phase column was used for the separation. Mobile phases were (A) 10 mM ammonium formate with 5 μM Agilent deactivator additive (Cat# 5191-3940) in 5:3:2 water:ACN:IPA and (B) 10 mM ammonium formate in 1:9:90 water:ACN:IPA. Column temperature was set at 55°C and autosampler temperature was at 20°C. The flow rate was 0.4 mL/min with sample injection volume of 4 µL. The following gradient was applied: 0 min, 15% B; 0-2.5 min, to 50% B; 2.5-2.6 min, to 57% B, 2.6-9 min, to 70% B; 9-9.1 min, to 93% B; 9.1-11.1 min, to 96% B; 11.1-15 min, 100% B; 15-20 min, 15% B.

Raw data were analyzed using Mass Hunter Qualitative analysis (10.0), MassProfinder 10.0 and MassProfiler Professional (MPP) 15.1 software (Agilent technologies). Lipid peak chromatograms were extracted against an in-house database created using MassHunter PCDL manager 8.0 (Agilent Technologies) and lipid standards, based on monoisotopic mass (< 5 ppm mass accuracy) and chromatographic retention time matches (< 0.5 min). Matched lipids were further confirmed by comparing MS/MS fragmentation pattern to the corresponding lipid species standards or publicly available MSMS databases (e.g. LIPID MAPS, HMDB, GNPS)

### Proteomics

#### Protein Sample Preparation and data analysis

Mouse muscle tissue was lysed in a buffer containing 50 mM ammonium bicarbonate (AMBIC), 4% SDS, and protease inhibitors. Samples were homogenized using an Omni TH tissue homogenizer, sonicated with a Branson sonicator, and clarified by centrifugation at 20,000 × g for 10 min. Protein concentration was determined, and 50 μg of total protein from each sample was subjected to TCA precipitation followed by an acetone wash. Pellets were re-suspended in 8□M urea, 50 mM AMBIC. Proteins were reduced and alkylated with dithiothreitol and iodoacetamide. Samples were diluted to 2□M urea with 50□mM AMBIC and digested overnight with Lysyl endopeptidase (lysC, Wako Chemicals USA, Inc.), then diluted to 1□M urea and digested with trypsin (Promega V5111) for 6 h. Peptides were desalted on C18 STAGE Tips [79]. Eluted peptides were dried and re-suspended in 5% formic acid.

The digests were analyzed using a Thermo Orbitrap Fusion mass spectrometer and an Easy nLC-1000 UHPLC. Peptides were separated with a gradient of 5–26% ACN in 0.1% FA over 125 min and introduced into the mass spectrometer by electrospray ionization as they eluted off a self-packed 40□cm, 100□µm (ID) column packed with 1.8 µm, 120 Å pore size, C18 resin (Sepax Technologies, Newark, DE). The column was heated to 60□°C. Peptides were detected using a data independent acquisition (DIA) mode. MS1 scans were collected in the Orbitrap mass analyzer overlap from 350-1200 m/z at 120K resolutions with the normalized automatic gain control (AGC) target set to 250 and a maximum ion accumulation time of 60 ms. The instrument was set to select precursors in 28 x 31.4 m/z wide windows with 1 m/z overlap for HCD fragmentation. The MS/MS scans were collected in the orbitrap at 30K resolution. The normalized AGC setting was 2000. Data were analyzed, searched, filtered, and assembled into protein quantification values using DIA-NN v1.8 [80] with the following settings: protease: Trypsin/P, missed cleavages: 1, peptide length range 7-30, FDR: 1%, match between runs: enable, quantification strategy: “Any LC (high accuracy)”, cross-run normalization: RT-dependent. The spectral library was generated in DIA-NN using *in silico* deep-learning and RT prediction based on the reviewed *Mouse* entries (downloaded March 6, 2021).

### Muscle homogenate and mitochondrial membranes preparation for enzymatic assays

Briefly, frozen muscle tissue was thinly cut on ice, and resuspended in isolation buffer containing 225 mM mannitol, 75 mM sucrose, 1 mM EGTA, 20 mM HEPES–KOH (pH 7.4), at tissue: buffer ratio 1:10 (w/v). Samples were homogenized for 1 min with a tissue-tearor IKA-type homogenizer (model 985370). Homogenates were centrifuged at 500 g for 5 min to remove debris and obtain the total tissue homogenate. To isolate crude mitochondrial membranes, the resulting homogenates were pelleted at 10,000 g for 10 min [81]. All procedures were performed at 4°C. Protein concentration was determined using bicinchoninic acid (BCA) assay.

### Complex I and Complex II activities

Enzyme activities were measured spectrophotometrically using a SpectraMax M5 plate reader (Molecular Devices) in 200 μl assay buffer essentially as described in [82]. Complex I NADH:HAR reductase activity was measured by monitoring NADH oxidation at 340 nm (ε340nm = 6.22 mM^-1^cm^-1^) in the presence of 1 mM hexaammineruthenium (HAR) as artificial electron acceptor. The experiments were performed in assay buffer (125 mM KCl, 20 mM HEPES-Tris (pH 7.4), 0.2 mM EGTA), supplemented with 40 µg/ml alamethicin, 1 mM KCN, and 1 mg/ml BSA. The reaction was initiated by the addition of 150 μM NADH, using 2-5 μg protein per well. Complex II succinate dehydrogenase activity was monitored following the reduction of dichlorophenolindophenol (DPCIP) at 600 nm (ε600nm = 21 mM^-1^cm^-1^) in assay buffer (20 mM HEPES-KOH, pH 7.8) containing 40 μg/ml alamethicin, 40 μM decylubiquinone (DBQ), 60 μM 2,6-dichlorophenol-indophenol (DCIP), 2 μM rotenone, and 0.5 mM KCN. The reaction was initiated by the addition of 20 mM succinate. Activity was fully inhibited by 5 nM atpenin A5.

### Complex V activity

To measure Complex V ATP-ase in gel activity, mitochondrial membranes were solubilized with N-dodecyl-β-D-maltoside (DDM) and subjected to high-resolution clear native (hrCN) electrophoresis as previously described [81]. ATP hydrolysis was measured in the presence of 6 mM lead (II) acetate, 14 mM MgCl_2_, and starting the reaction with 8 mM ATP in a buffer composed of 30 mM Tris and 270 mM glycine (pH 8.5 A total of 50 μg of protein was loaded per lane. The gel was incubated overnight at room temperature, and densitogram analysis was performed using a LI-COR Odyssey CLx imaging system (ImageStudio software).

### Statistical analysis

In all assays the values are averages of at least three independent measurements. Error bars indicate standard deviation (SD) or standard error of the mean (SEM). Statistically significant differences between two groups were estimated by unpaired two-tailed Student’s test with significance set at p<0.05. Biological replicates information for each experiment is indicated in the figure legends. Lipid peak areas and median-normalized protein abundance were used for lipidomics and proteomics data analyses, respectively.

## AUTHOR CONTRIBUTIONS

**Inaya Laubach**: Data curation; formal analysis; investigation; writing – review and editing. **Guido Primiano**: Data curation; formal analysis; investigation; writing – review and editing. **Nneka Southwell**: Data curation; formal analysis; investigation; writing – review and editing. **Nicola Rizzardi**: Data curation; formal analysis; investigation; writing – review and editing. **Belem Yoval-Sánchez**: Data curation; formal analysis; methodology; writing – review and editing. **Christian Bergamini**: Conceptualization; supervision. **Serenella Servidei**: Conceptualization; supervision. **Alexander Galkin**: Conceptualization; formal analysis; funding acquisition; writing – review and editing. **Giovanni Manfredi**: Conceptualization; formal analysis; supervision; funding acquisition; writing– review and editing. **Qiuying Chen**: Conceptualization; formal analysis; supervision; funding acquisition; validation; writing – original draft; writing – review and editing. **Marilena D’Aurelio**: Conceptualization; formal analysis; supervision; funding acquisition; validation; writing – original draft; writing – review and editing.

## FOR MORE INFORMATION

Supporting information includes four figures and one table.

## ACKNOWLEDGMENTS

We thank Dr. Zhucui Li at Weill Cornell Proteomics and Metabolomics Core Facility, the Weill Cornell Metabolic Phenotyping Center, the Weill Cornell Microscopy and the Image Analysis Core for their contribution. Schematics were created with Biorender.com

