## Supplemental information for "Lipid mediated ER-stress contributes to the pathogenesis of mitochondrial myopathies"

### FIG. S1

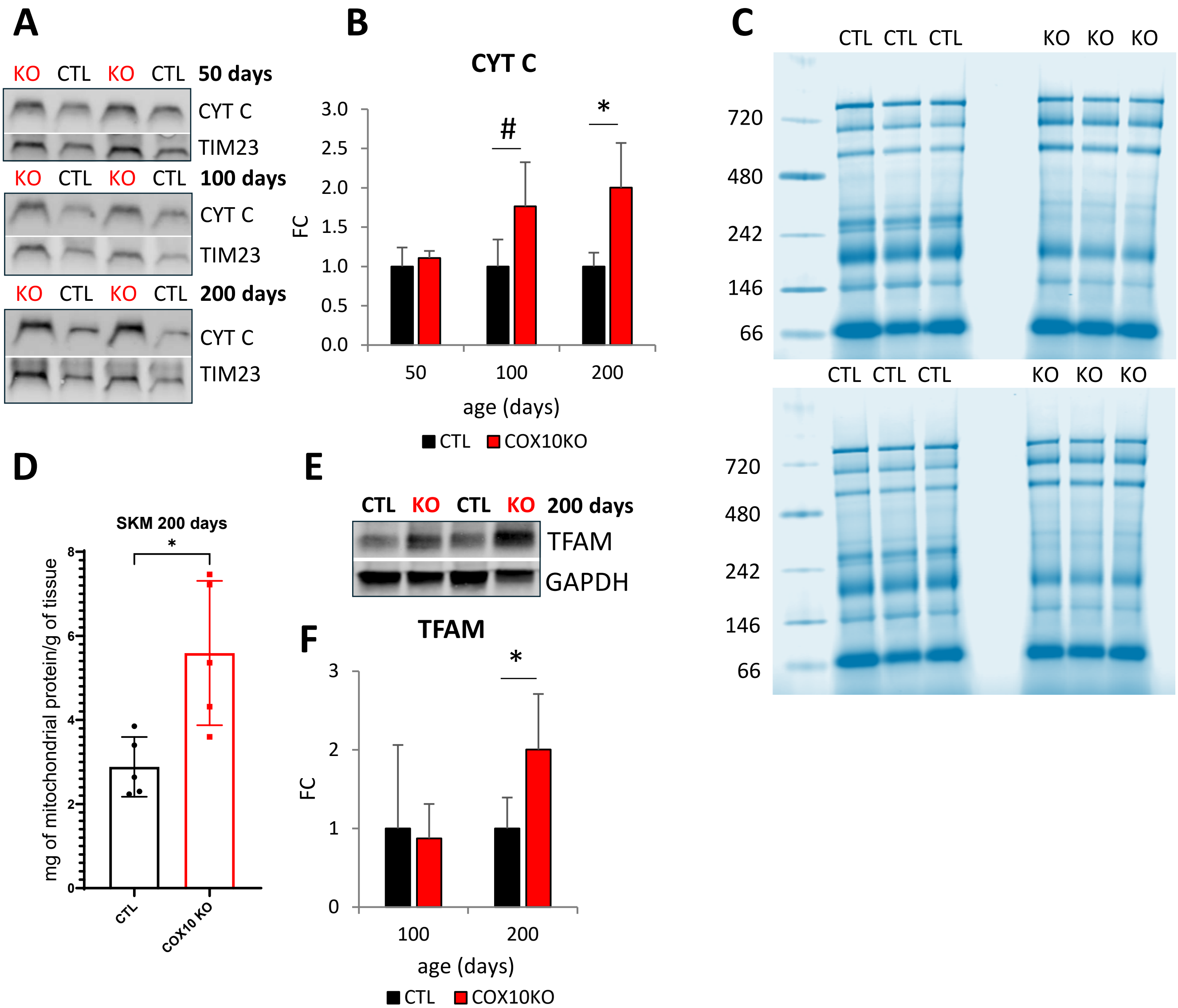


**FIGURE S1**

**A)** Representative WB of muscle lysates from 50, 100, and 200 days old COX10 KO and CTL mice, separated by denaturing SDS-PAGE, and probed for Cytochrome c and TIM23.
**B)** Age-dependent protein levels of Cytochrome c in COX10 KO muscle (n=4 per age group), normalized by TIM23, expressed relative to CTL (n=4 per age group), at 50, 100, and 200 days. Data are presented as Mean ± SD, * $p<0.05$, # $p=0.06$, COX10 KO vs. same age CTL set at 1, by unpaired t-test.
**C)** Comassie blue staining of BN-PAGE prior to transfer of samples in Fig 1I. The prominent band at 66 kDa across all lanes indicates similar sample loading.
**D)** Mitochondrial content in skeletal muscle (SKM) of 200 days old COX10 KO (n=5) and CTL (n=5) mice, assessed as mitochondrial protein per gram of tissue.
**E)** Representative WB of muscle lysates from 200 days old COX10 KO and CTL mice, separated by denaturing SDS-PAGE, and probed for TFAM and GAPDH.
**F)** Age-dependent protein levels of TFAM in COX10 KO muscle (n=4 per age group), normalized by GAPDH, expressed relative to CTL (n=4 per age group), at 100 and 200 days. Data are presented as Mean ± SD, * $p<0.05$ COX10 KO vs. same age CTL set at 1, by unpaired t-test.

### FIG. S2

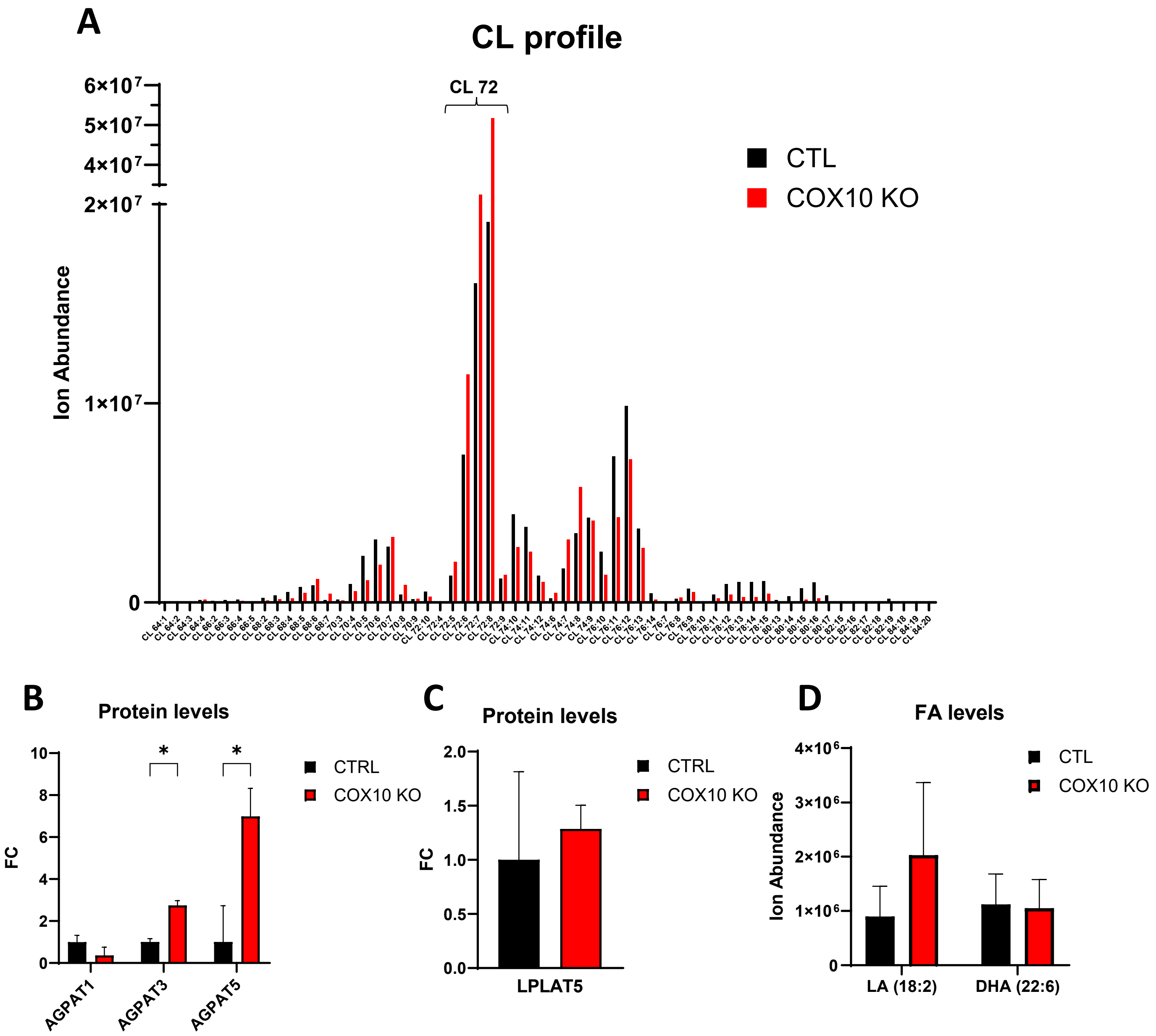


**FIGURE S2**

**A)** Muscle CL profile of COX10 KO (n=4) and CTL (n=4) mice, measured by LC/MS analysis, at 150 days. Data are presented as Mean.

**B)** Muscle protein levels of AGPAT1, AGPAT3, and AGPAT5 in COX10 KO (n=4) expressed relative to CTL (n=4) set at 1, by proteomics analysis, at 200 days. Data are presented as Mean ± SD, * $p<0.05$, by unpaired t-test.

**C)** Muscle protein levels of LPLAT5 in COX10 KO (n=4) expressed relative to CTL (n=4) set at 1, by proteomics analysis, at 200 days. Data are presented as Mean ± SD.

**D)** Muscle FA levels in COX10 KO (n=4) and CTL (n=4) mice, measured by LC/MS analysis, at 150 days. Data are presented as Mean ± SD.

### FIG. S3

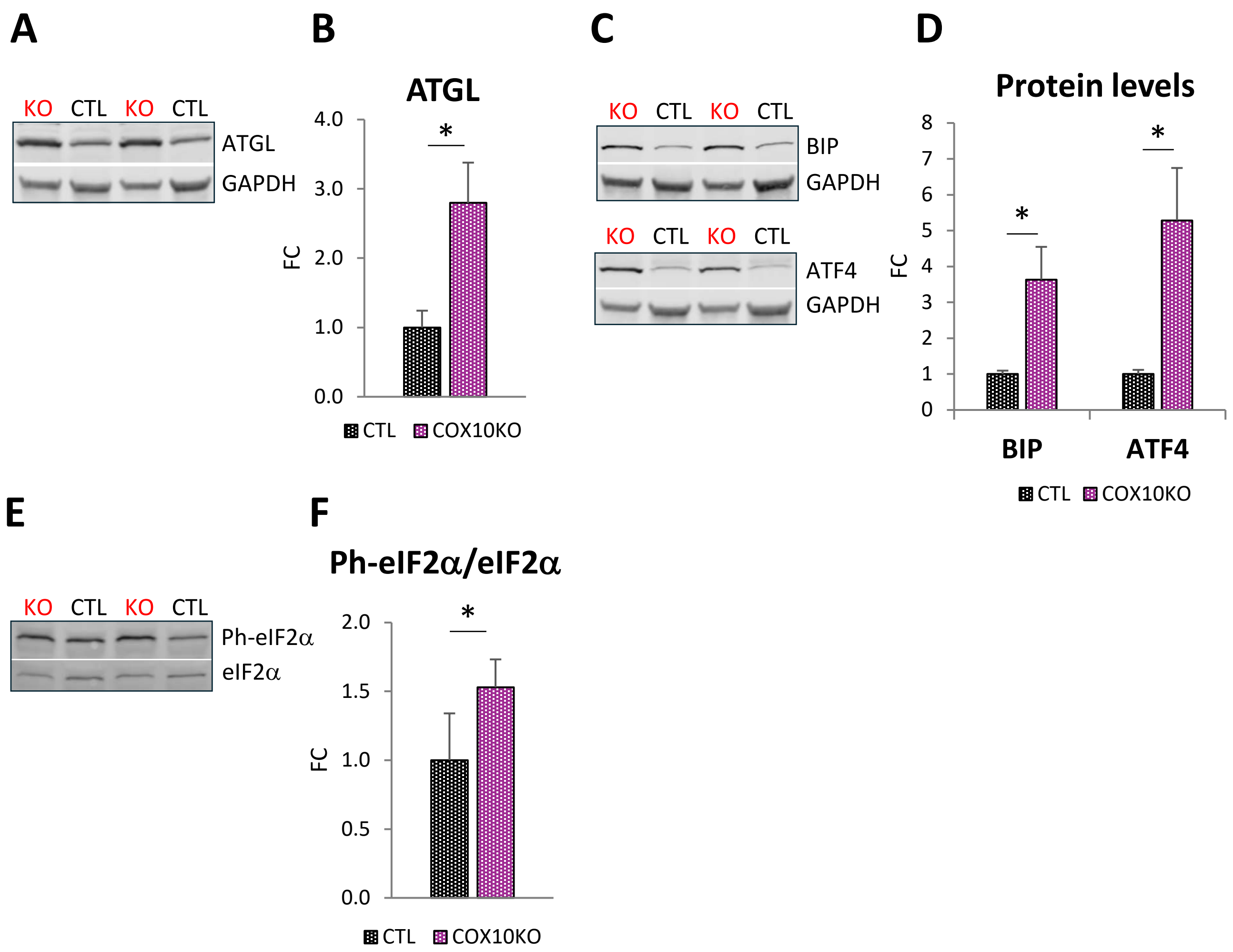


**FIGURE S3**

**A)** Representative WB of muscle lysates from 200 days old COX10 KO and CTL female mice, separated by denaturing SDS-PAGE, and probed for ATGL and GAPDH.
**B)** Muscle protein levels of ATGL in 200 days old female COX10 KO (n=4), normalized by GAPDH, expressed relative to 200 days old female CTL (n=4) set at 1. Data are presented as Mean ± SD, * $p<0.05$, by unpaired t-test.
**C)** Representative WB of muscle lysates from 200 days old COX10 KO and CTL female mice, separated by denaturing SDS-PAGE, and probed for BIP, ATF4, and GAPDH.
**D)** Muscle protein levels of BIP and ATF4 in 200 days old female COX10 KO (n=4), normalized by GAPDH, expressed relative to 200 days old female CTL (n=4) set at 1. Data are presented as Mean ± SD, * $p<0.05$, by unpaired t-test.
**E)** Representative WB of muscle lysates from 200 days old COX10 KO and CTL female mice, separated by denaturing SDS-PAGE, and probed Ph-eIF2α and eIF2α.
**F)** Muscle levels of Ph-eIF2α/eIF2α in 200 days old female COX10 KO (n=4) expressed relative to 200 days old female CTL (n=4) set at 1. Data are presented as Mean ± SD, * $p<0.05$, by unpaired t-test.

FIG. S4

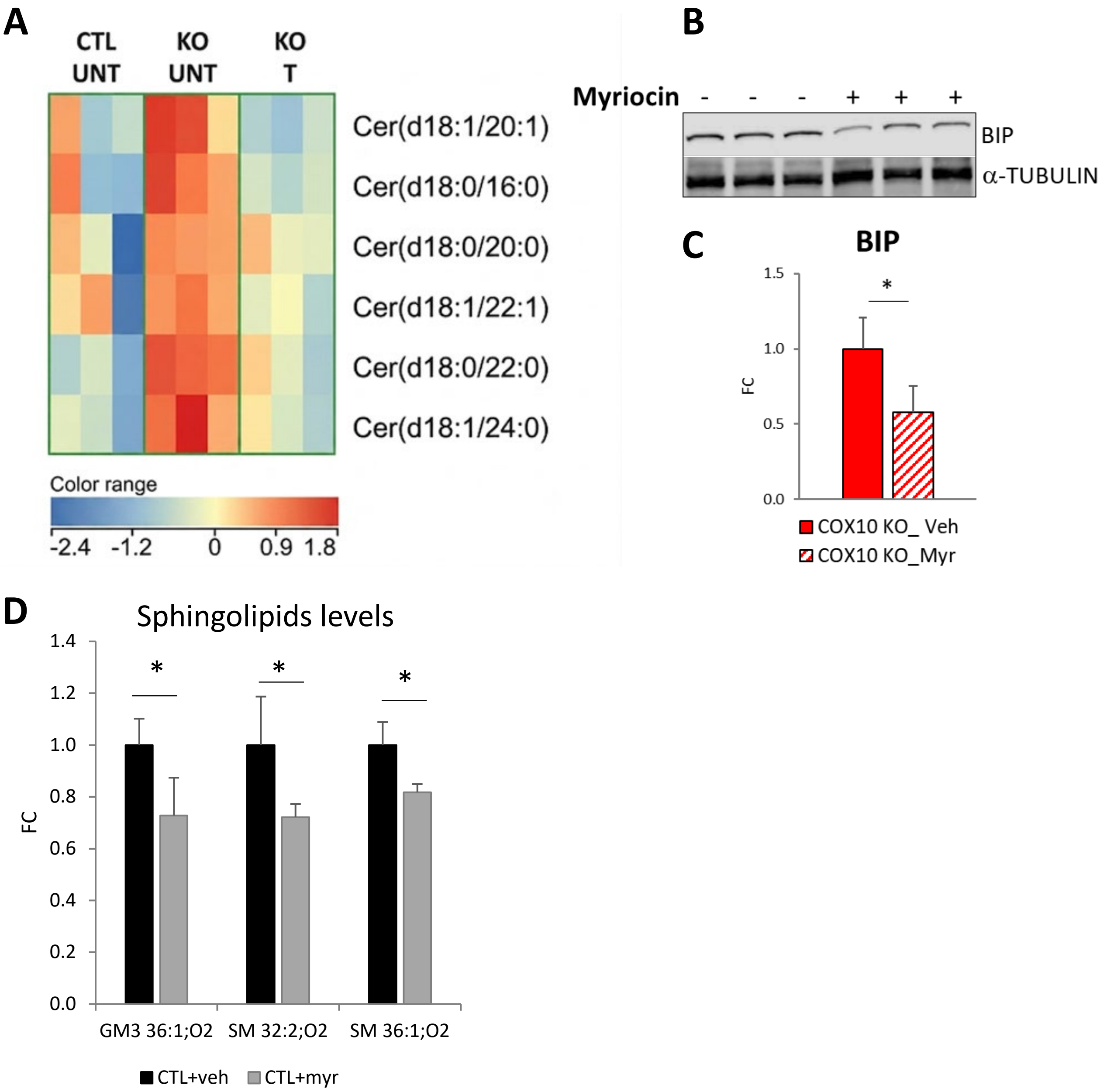


**FIGURE S4**

**A)** Muscle Ceramide (Cer, d18:1) and dihydroceramide (DH-Cer, d18:0) by LC/MS analysis in COX10 KO treated for 10 days with myriocin (T) and COX10 KO and CTL mice treated with vehicle (UNT), n=3 per group, at 130 days. Colors are based on the Z-score, blue denotes lower abundance and red higher abundance.

**B)** WB of muscle lysates from 130 days old COX10 KO mice treated with myriocin or vehicle, separated by denaturing SDS-PAGE, and probed for BIP and α-TUBULIN.

**C)** Muscle protein levels of BIP in COX10 KO treated with myriocin (n=6) normalized by α-TUBULIN, expressed relative to COX10 KO treated with vehicle (n=3) set at 1. Data are presented as Mean ± SD, * $p<0.05$, by unpaired t-test.

**D)** Muscle levels of a subset of sphingolipids in CTL mice treated with myriocin (n=4) and vehicle (n=4) for 3 months, by LC/MS analysis, at 150 days. Data are presented as Mean ± SD, * $p<0.05$, CTL with myriocin vs. CTL with vehicle set at 1, by unpaired t-test.

### Supporting Info Table

#### Table S1: Patient and CTL information.

| Diagnosis | Specimen | Sex | Age at diagnosis | Mutation | COX-negative fibers | % Muscle Heteroplasmy |
|---|---|---|---|---|---|---|
| MERRF | Muscle (4) | F(1), M(3) | 44±6 | m.8344A>G/*MT-TK* | + | 82±5 |
| Healthy CTL | Muscle (4) | F(2), M(2) | 45±4 | N/A | - | N/A |

Data are presented as Mean ± SD. F= females, M= Males. The number of individuals in each group is indicated in parenthesis. N/A=data not available. Presence (+) or absence (-) of COX-negative fibers.
